# Arachidonic acid incorporation into phosphatidylinositol by LPLAT11/MBOAT7 ensures radial glial cell integrity in developing neocortex

**DOI:** 10.1101/2024.04.30.588048

**Authors:** Yuki Ishino, Yusuke Kishi, Taiga Iwama, Naohiro Kuwayama, Hiroyuki Arai, Yukiko Gotoh, Junken Aoki, Nozomu Kono

## Abstract

Arachidonic acid, a vital polyunsaturated fatty acid in brain development, is enriched in phosphatidylinositol (PI). The arachidonic acyl chain in PI is introduced by lysophospholipid acyltransferase 11 (LPLAT11)/membrane-bound O-acyltransferase 7 (MBOAT7), the loss of which causes cortical atrophy in humans and mice. Here, we show that LPLAT11 deficiency impaired indirect neurogenesis in the developing neocortex, resulting in fewer layer II-V neurons. LPLAT11-deficient radial glial cells had defects in differentiation into intermediate progenitor cells and increased apoptosis. Prior to these anomalies, LPLAT11 deficiency caused a fragmentation of the Golgi apparatus, accompanied by impaired apical trafficking of E-cadherin, and deregulated apical detachment. Moreover, impaired PI acyl chain remodeling led to a decreased amount of PI(4,5)P_2_, leading to Golgi apparatus fragmentation. Thus, these results clarify the underlying mechanism of cortical atrophy by LPLAT11 deficiency and highlight the critical role of arachidonic acid in PI in the integrity of radial glial cells.

## Introduction

Arachidonic acid is a major polyunsaturated fatty acid (PUFA) found in the brain and plays a crucial role in neuronal development and function^1,2^. Classically, its physiological impact is attributed to its conversion into prostaglandins and leukotrienes; however, the functions of arachidonic acid-derived mediators only partially explain its significance in brain development^3–6^. Arachidonic acid is primarily present as a phospholipid-bound state and is particularly enriched in the *sn*-2 position of phosphatidylinositol (PI). The enrichment of arachidonic acid in PI occurs through the remodeling of the acyl chain at the *sn*-2 position, at which newly synthesized PI undergoes deacylation and re-acylation^7–10^. Previously, we identified lysophospholipid acyltransferase 11 (LPLAT11, also known as LPIAT1 or MBOAT7) as the remodeling enzyme that specifically incorporates arachidonic acid into PI^11^. Recent research has revealed that patients harboring null mutations in *MBOAT7*, which encodes LPLAT11, show severe intellectual disability, epilepsy, autistic features, and cortical atrophy^12–20^. Consistent with this, *Mboat7* knockout (KO) mice also showed the atrophy of the cerebral cortex at embryonic day 18.5 (E18.5)^21,22^. Given that arachidonic acid-derived mediators are not affected in the cortex of *Mboat7* KO mice^21^, it is likely that a decrease in arachidonic acid-bound phosphatidylinositol, rather than a decrease in the mediators, is responsible for cortical atrophy. However, the underlying mechanism linking LPLAT11 deficiency to brain malformation remains unclear.

Cortical atrophy results from fewer neurons generated during the embryonic stages. In the developing mammalian cerebral cortex, neural stem cells (neuroepithelial cells) undergo self-renewal through symmetric divisions until around E10. At the early stage of neurogenesis (E11-E13), neural stem cells (radial glial cells, RGCs) generate neurons mainly by asymmetric divisions in a process called direct neurogenesis. During mid-corticogenesis (E13-E16), RGCs generate neurons not only by direct neurogenesis but also by indirect neurogenesis, in which RGCs spawn intermediate progenitor cells (IPCs) mainly through asymmetric divisions and IPCs generate neurons after 1-2 times of cell division^23–26^. Whereas direct neurogenesis generates mostly deep-layer neurons, indirect neurogenesis can generate all layers of neurons, especially upper-layer neurons^27,28^. Therefore, inhibition of indirect neurogenesis reduces the number of neurons and causes cortical atrophy^29^.

In the present study, we showed that LPLAT11 deficiency compromised the integrity of RGCs in the E12 cortex. LPLAT11-deficient RGCs exhibited impaired indirect neurogenesis and increased apoptosis, leading to microcephaly as a result of fewer layer II-V neurons. Prior to these anomalies, we observed a fragmented Golgi apparatus, reduced expression of E-cadherin on the apical surface, and apical detachment, which, together, suggest trafficking of E-cadherin to the apical surface is impaired. Furthermore, LPLAT11 deficiency reduced the amount of PI(4,5)P_2_, leading to fragmentation of the Golgi apparatus. These findings underscore the critical role of arachidonic acid in PI in the integrity of RGCs.

## Results

### Layer II-V neurons decrease in LPLAT11-deficient mice

The cerebral cortex comprises six neural layers. To investigate which cortical layers were affected by LPLAT11 deficiency, we immunostained E18.5 cortices using antibodies against T-box brain 1 (Tbr1), COUP-TF interacting protein 2 (Ctip2), and Cut-like homeobox 1 (Cux1), which are markers for layer VI/subplate, layer VI/V, and layer II-IV neurons, respectively (Fig. 1a, b). Although the number of layer VI neurons in *Mboat7* KO mice was comparable to those in *Mboat7* heterozygous mice, the numbers of layer V neurons and layer II-IV neurons decreased to 70% and 50% of those in *Mboat7* heterozygous mice, respectively (Fig. 1a-c). Neurons in layer VI are generated from E12.5 RGCs, whereas neurons in layers II-V are generated from E13.5-E16.5 RGCs^30^. These results suggest that LPLAT11 deficiency impairs neuron production during E13.5-E16.5.

**Fig. 1.**
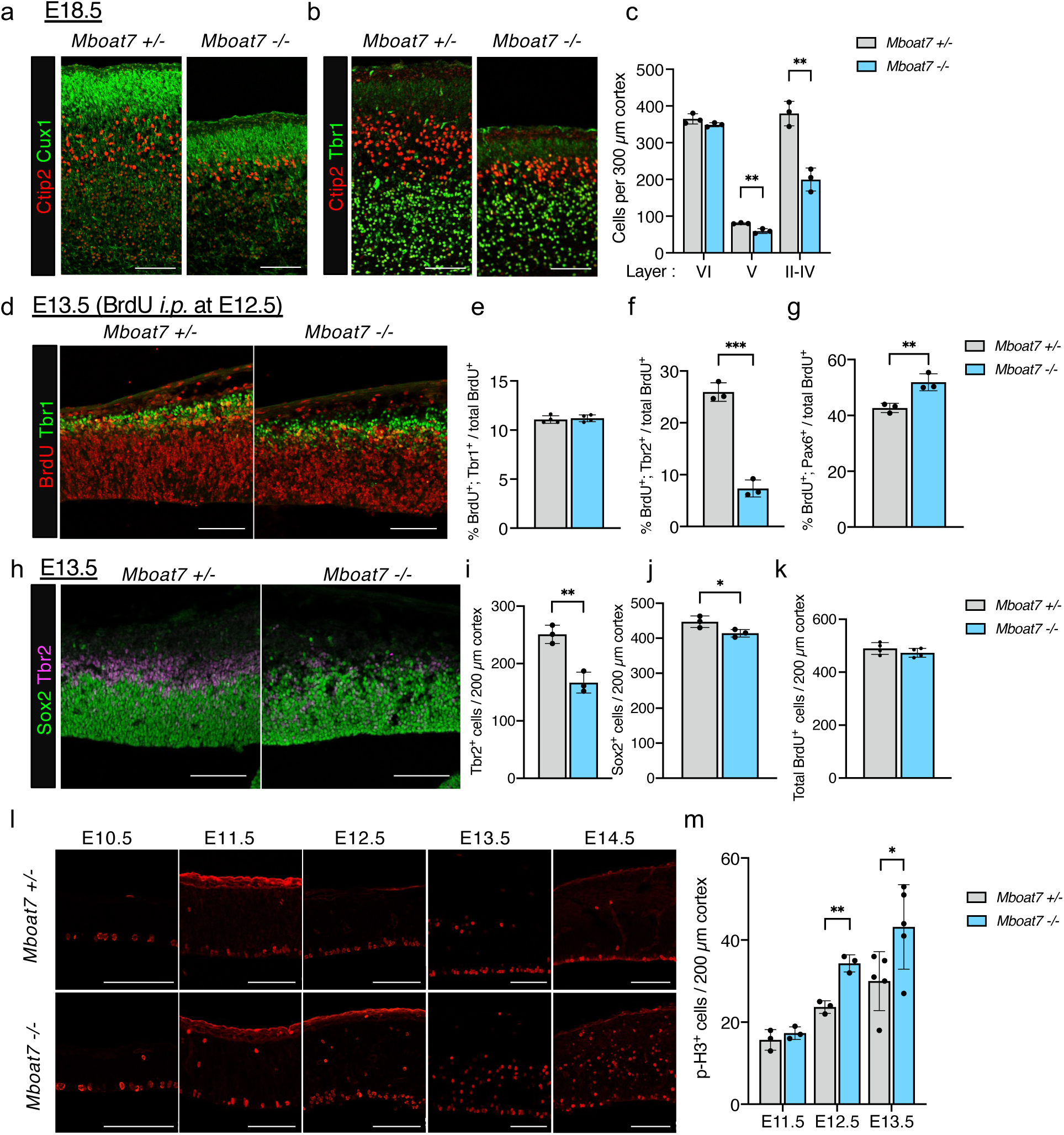
LPLAT11-deficient mice show the decrease in the number of layer II-V neurons, the inhibition of the differentiation into IPCs, and aberrant progenitor mitosis. (a) Immunofluorescence staining of Cux1 (green) and Ctip2 (red) in coronal sections of *Mboat7* heterozygous and KO mice at E18.5. (b) Immunofluorescence staining of Tbr1 (green) and Ctip2 (red) in coronal sections of *Mboat7* heterozygous and KO mice at E18.5. (c) Quantitative analysis of layer VI (Tbr1^+^), V (Tbr1^-^; Ctip2^+^), and II-IV (Cux1^+^) neurons per area within 300 µm wide bins (n=3 mice for each genotype). (d) Double staining at E13.5 for BrdU (injected at E12.5) and Tbr1. Arrows show neurons (BrdU^+^; Tbr1^+^). (e) The rate of neuronal differentiation (ratio of BrdU^+^; Tbr1^+^ /BrdU^+^ cells) (n=4 mice for each genotype). (f) The rate of differentiation into IPCs (ratio of BrdU^+^; Tbr2^+^ /BrdU^+^ cells) (n=3 mice for each genotype). (g) The rate of RGCs (ratio of BrdU^+^; Pax6^+^ /BrdU^+^ cells) (n=3 mice for each genotype). (h) Immunofluorescence staining of Sox2 (green) and Tbr2 (magenta) in coronal sections of *Mboat7* heterozygous and KO mice at E13.5. (i,j) Quantitative analysis of cells positive for Tbr2 (i) and Sox2 (j) per area within 200 µm wide bins (n=3 mice for each genotype). (k) Total BrdU^+^ cells in 200 µm cortex at E13.5 (BrdU injected at E12.5) (n=4 mice for each genotype). (l) Immunofluorescence staining for phospho-histone H3 (p-H3) in the cortices of *Mboat7* heterozygous and KO mice from E10.5 to E14.5. (m) Quantitative analysis of cells positive for p-H3 per area within 200 µm wide bins (n=3-5 mice for each genotype). Data are shown as mean ± SEM. Unpaired two-tailed Student’s t-test; *p<0.05, **p<0.01 and ***p<0.001. Scale bars, 100 µm.

### Differentiation into IPCs from RGCs is impaired in LPLAT11-deficient mice

The decrease in the number of neurons can be caused by several factors, such as perturbation of neuron differentiation from RGCs, apoptosis of neurons/neuron-producing cells, and decreased proliferation of RGCs. We first examined whether neuron differentiation is inhibited in *Mboat7* KO mice. E13.5 pregnant mice were intraperitoneally injected with bromodeoxyuridine (BrdU), and the E14.5 embryo brain was analyzed. Double staining for Tbr1 and BrdU revealed that Tbr1^+^; BrdU^+^ cells were comparable between *Mboat7* heterozygous and KO mice (Fig. 1d,e), showing that neuronal differentiation was normal in *Mboat7* KO mice. Triple staining for Pax6 (RGC marker), Tbr2 (IPC marker), and BrdU revealed that Tbr2^+^; BrdU^+^ cells decreased, whereas Pax6^+^; BrdU cells increased in *Mboat7* KO mice (Fig. 1f,g). We also quantified Tbr2-and Sox2 (RGC marker)-positive cells irrespective of BrdU labeling. Tbr2^+^ cells also decreased in *Mboat7* KO mice, whereas the number of Sox2^+^ cells was slightly decreased (Fig. 1h-j). These results indicate that the differentiation into IPCs is inhibited in *Mboat7* KO mice.

### Phosphorylation of c-Jun increases in RGCs in LPLAT11-deficient mice

Tbr2 expression is inhibited by c-Jun activation in immune cells^31^. To investigate whether c-Jun phosphorylation is increased in the *Mboat7* KO cortex, E10.5-E14.5 cortices were immunostained or immunoblotted for phospho-c-Jun (p-c-Jun). Phosphorylation of c-Jun was increased from E12.5 and peaked at E14.5 in the cortices of *Mboat7* KO mice (Supplementary Fig. 1a-c). The protein levels of c-Jun N-terminal Kinase (JNK) and its phosphorylated form (p-JNK) were not changed in the cortices of *Mboat7* KO mice (Supplementary Fig. 1d,e), indicating that increased c-Jun phosphorylation was independent of JNK. Double staining for p-c-Jun and cell type-specific markers revealed that c-Jun phosphorylation was upregulated in the cell population below the cortical plate (Supplementary Fig. 1f) and p-c-Jun merged with Sox2 but not with Tbr2 (Supplementary Fig. 1g,h), indicating that phosphorylation of c-Jun was upregulated specifically in *Mboat7* KO RGCs.

### Proliferation of RGCs is not decreased in LPLAT11-deficient mice

We next evaluated the proliferation of neural progenitors. BrdU labeling experiments showed that total BrdU^+^ cells were comparable between *Mboat7* heterozygous and KO mice (Fig. 1k). However, immunostaining for phospho-Histone H3 (p-H3), a marker for M-phase cells, in E10.5-E14.5 cortices showed that the number of p-H3-positive cells started to increase at E12.5 in the cortices of *Mboat7* KO mice compared to *Mboat7* heterozygous mice (Fig. 1l,m), suggesting that the proliferation of neural progenitors is not impaired in *Mboat7* KO mice, although their mitosis is aberrant.

### Apoptosis is increased specifically during E13.5-E15.5 in the cortices of LPLAT11-deficient mice

To examine apoptosis in the cortices of *Mboat7* KO mice, we collected cortices from E12.5 to E18.5 embryos and evaluated caspase-3 activation by western blot analysis. We found that apoptosis was markedly increased during the period of E13.5-E15.5 in the cortices of *Mboat7* KO mice (Fig. 2a,b). Furthermore, immunostaining for cleaved caspase-3 in E10.5-E14.5 cortices revealed that apoptosis was not clearly observed until E12.5 in *Mboat7* KO mice (Fig. 2c,d). Co-immunostaining of *Mboat7* KO cortices with cleaved caspase-3 and Tbr2, Ctip2, or Sox2 showed that neither Tbr2, Ctip2, nor Sox2 merged with cleaved caspase-3 (Fig. 2e-g), although cleaved caspase-3-positive cells localized exclusively in the ventricular and subventricular zones.

**Fig. 2.**
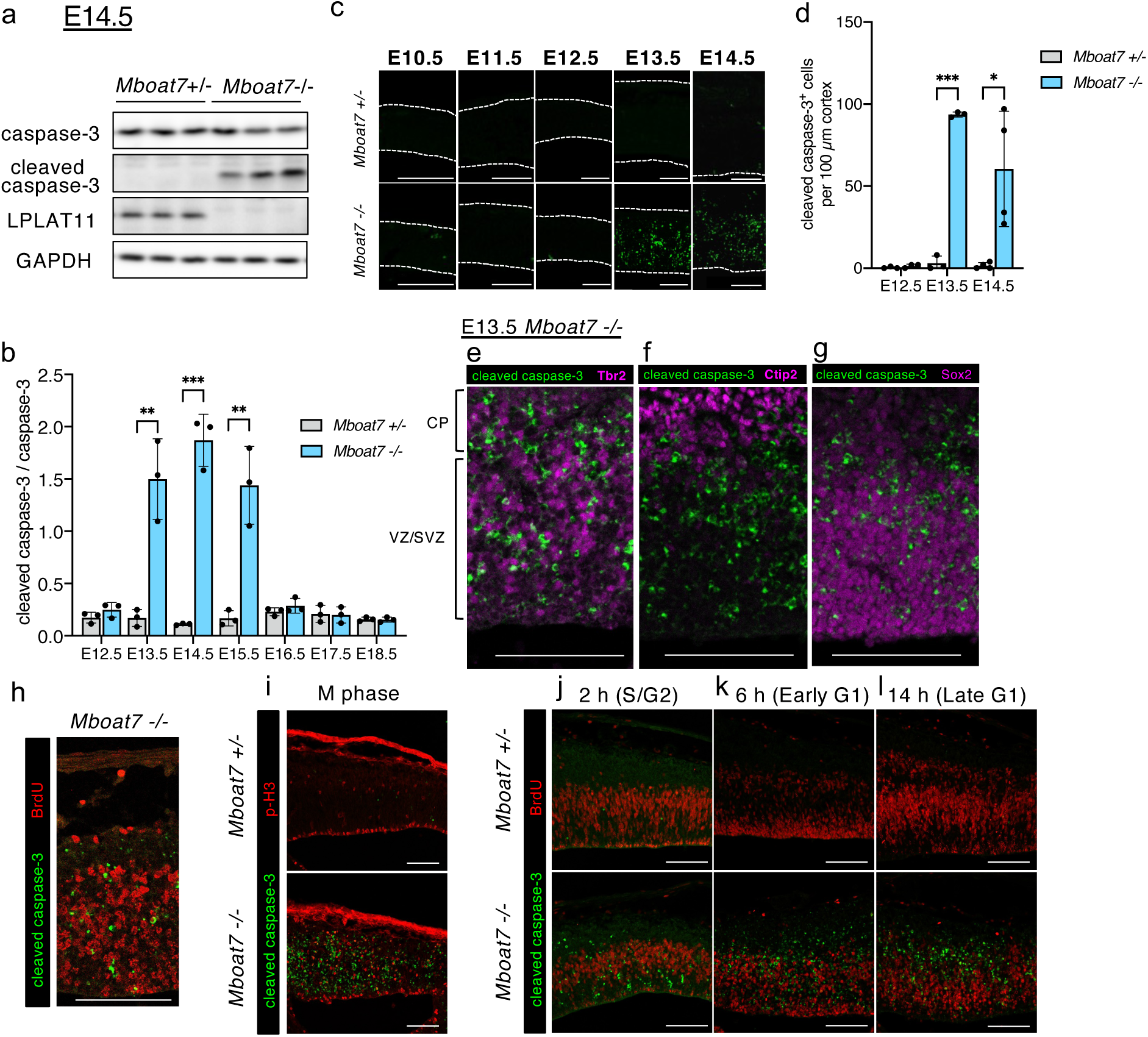
LPLAT11-deficient mice show increased apoptosis in the developing cortex. (a) Western blots of caspase-3, cleaved caspase-3 and LPLAT11 in the cortex of *Mboat7* heterozygous and KO mice at E14.5. GAPDH is used as a loading control. (b) Evaluation of caspase-3 activation in the cortex of *Mboat7* heterozygous and KO mice from E12.5 to E18.5 (n=3 mice for each genotype). (c) Immunofluorescence staining for cleaved caspase-3 in the cortex of *Mboat7* heterozygous and KO mice from E10.5 to E14.5. Dotted lines show the apical and the basal surfaces of the cortex. (d) Quantitative analysis of cells positive for cleaved caspase-3 per area within 100 µm wide bins (n=3-4 mice for each genotype). (e-h) Double staining for cleaved caspase-3 and Tbr2 (e), Ctip2 (f), Sox2 (g), or BrdU (h) in *Mboat7* KO mice cortex at E13.5. (i) Double staining for cleaved caspase-3 and p-H3 in the E13.5 cortices of *Mboat7* heterozygous and KO mice. (j-l) Double staining for cleaved caspase-3 and BrdU in the E13.5 cortices of *Mboat7* heterozygous and KO mice, labeled for 2 h (j), 6 h (k) and 14 h (l) in BrdU labeling. CP, cortical plate; VZ, ventricular zone; SVZ, subventricular zone. Data are shown as mean ± SEM. Unpaired two-tailed Student’s t-test; *p<0.05, **p<0.01 and ***p<0.001. Scale bars,100 µm.

We noticed that cleaved caspase-3-positive cells didn’t merge with BrdU (Fig. 2h). Thereby, we further examined at which phase of the cell cycle cells underwent apoptosis in *Mboat7* KO mice. To label proliferating cells in the S/G2 phases, early G1 phase, and late G1 phase, we injected BrdU at 2 h, 6 h, and 14 h, respectively, before harvesting at E13.5, and then E13.5 cortices were co-immunostained with cleaved caspase-3 and BrdU. In any case, cleaved caspase-3 didn’t merge with BrdU staining, indicating that cells with defects in the entry into the S phase undergo apoptosis (Fig. 2j-l). Moreover, cleaved caspase-3-positive cells hardly merged with p-H3, suggesting that M-phase cells were not the major cells that underwent apoptosis (Fig. 2i). Taken together, these results suggest that mitotic arrest followed by apoptosis (which is known as mitotic catastrophe) occurs in the *Mboat7* KO cortex.

### Cyclin-dependent kinases are involved in the apoptosis in LPLAT11-deficeint mice

We focused on cyclin-dependent kinases (CDKs) because CDKs are reported to be involved in mitotic catastrophe^32^ and are also one of the kinases of c-Jun phosphorylation^33,34^. Indirubin 3’-monoxime (I3M), a potent CDK inhibitor, was injected into the lateral ventricle at E11.5, followed by immunostaining for p-H3 in the cortex at E12.5. I3M administration decreased the signal of p-c-Jun at the ventricular surface and the number of p-H3-positive RGCs at the ventricular zone (Supplementary Fig. 2a).

To examine whether CDK inhibition suppresses apoptosis in *Mboat7* KO mice, we used a cortical hemisphere culture because this culture system has been reported to approximate *in vivo* and be useful for assessing the effect of pharmacological treatment^35^. In the hemisphere culture system, the E12.5 hemispheres were cultured in a shaker for 24 h and collected (Supplementary Fig. 2b). We observed apoptosis in the *Mboat7* KO E12.5 hemisphere cultured for 24 h and I3M treatment suppressed the apoptosis in the ventricular zone (Supplementary Fig. 2c). These results indicate that aberrant CDK activation is the cause of mitotic catastrophe in *Mboat7* KO mice.

### E-cadherin expression decreases in the apical surface of LPLAT11-deficient mice

RGCs constantly extend their processes radially and adhere to the apical and basal surfaces of the cortex. Nuclei of RGCs migrate basally during the G1/S phase and apically during the G2/M phase^36^. Therefore, RGCs divide at the apical surface. We noticed that M-phase cells dispersed within ventricular/subventricular zone in *Mboat7* KO mice starting at E12.5, whereas M-phase cells localized to the apical surface in *Mboat7* heterozygous mice (Fig. 1l). In the E13.5 cortex of *Mboat7* KO mice, RGCs and IPCs dispersed within the ventricular zone/subventricular zone (Fig. 3a,b and Supplementary Fig. 3a,b), whereas Ctip2-positive neurons normally localized to the cortical plate (Supplementary Fig. 3c,d). To distinguish M-phase RGCs from IPCs, E13.5 cortices of *Mboat7* KO mice were immunostained for Pax6 (or Sox2), Tbr2, and p-H3. Although some M-phase RGCs (p-H3^+^; Pax6^+^) localized at the apical surface, others did not localize to the apical surface in *Mboat7* KO mice (Fig. 3a-c), suggesting that *Mboat7* KO RGCs lost their apical processes. We also found some unusual cells that are negative for both Sox2 and Tbr2 and a few immature IPCs (Sox2^+^; Tbr2^+^) in p-H3-positive cells in *Mboat7* KO cortices. To examine the apical and basal processes of RGCs in *Mboat7* KO mice, E12.5 and E13.5 cortices were stained with Nestin. In E13.5 cortices, the processes of RGCs extended radially from the apical surface to the basal surface, and Nestin was strongly stained at the apical surface in *Mboat7* heterozygous mice (Fig. 3d). On the contrary, in *Mboat7* KO mice, Nestin was weakly stained at the apical surface, and the radial fiber arrangement was disordered and showed a honeycomb-like pattern in *Mboat7* KO mice, suggesting that RGCs lost contact at the apical surface and detached from the apical surface in *Mboat7* KO mice cortex. The disordered arrangement of radial fibers was also observed in E12.5 cortices of *Mboat7* KO mice (Fig. 3d, inset).

**Fig. 3.**
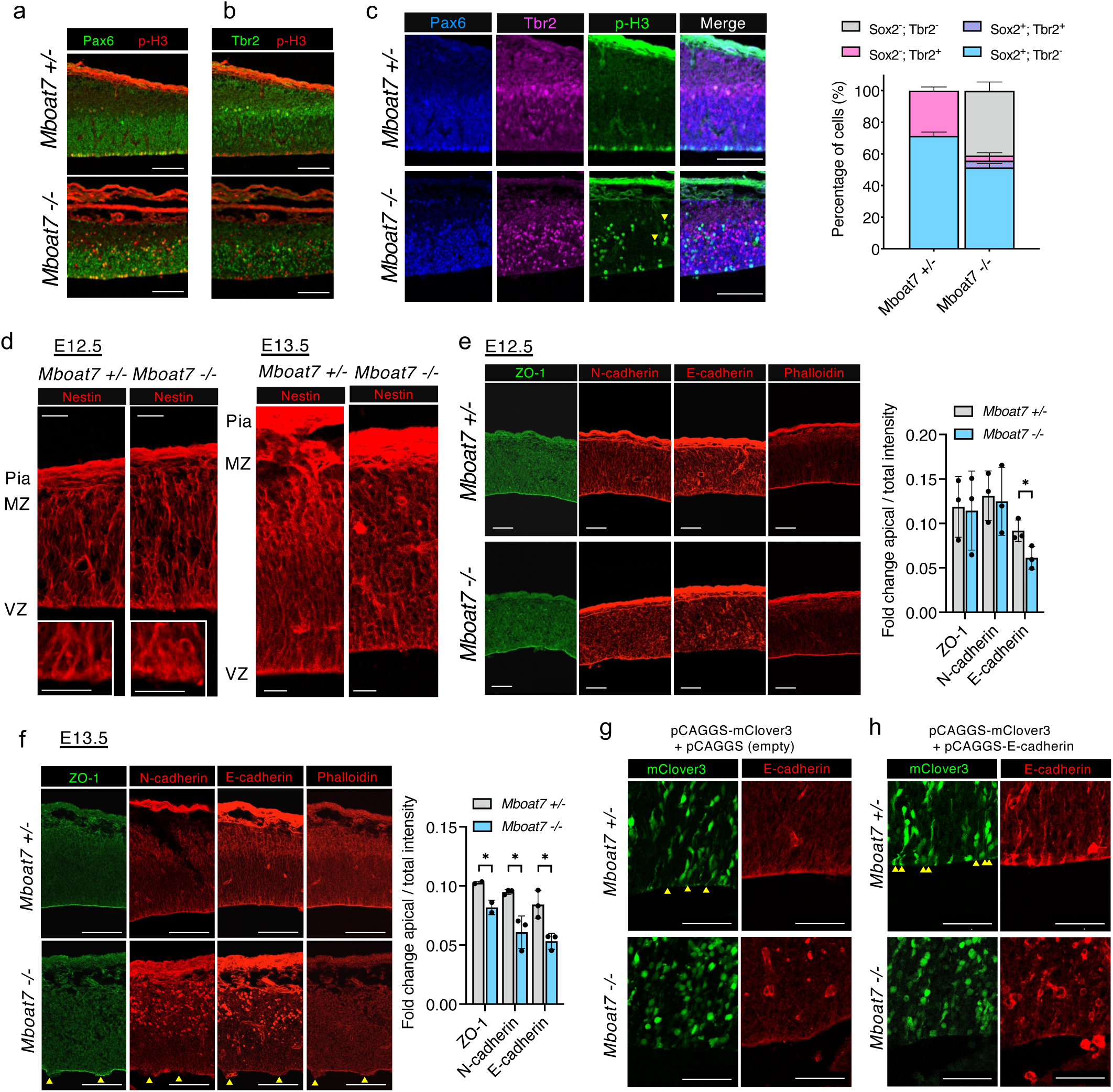
Adherens junction is impaired in the cortex of LPLAT11-deficient mice. (a,b) Double staining for Pax6 and p-H3 (a) or p-H3 and Tbr2 (b) in the cortices of *Mboat7* heterozygous and KO mice at E13.5. (c) Triple staining for Pax6, Tbr2 and p-H3 in the cortices of *Mboat7* heterozygous and KO mice at E13.5. Ratio of Pax6^+^; Tbr2^-^ RGCs (cyan), Pax6^+^; Tbr2^+^ immature IPCs (purple), Pax6^-^; Tbr2^+^ IPCs (magenta) and Pax6^-^; Tbr2^-^ abnormal cells (grey) were shown in stacked bar charts (n=3 mice for each genotype). (d) Immunofluorescence staining for Nestin in the cortices of *Mboat7* heterozygous and KO mice at E13.5. Pia, pial surface; MZ, marginal zone; VZ, ventricular zone. (e,f) Immunofluorescence staining for ZO-1, N-cadherin, E-cadherin, and phalloidin in the cortices of *Mboat7* heterozygous and KO mice at E12.5 (e) and E13.5 (f). Fold change of apical intensity against total intensity from VZ to MZ is shown for ZO-1, N-cadherin and E-cadherin (n=2,3 mice for each genotype). Arrowheads show the disruption of the ventricular wall. (g,h) pCAGGS-mClover3 and pCAGGS empty vector (g) or pCAGGS-E-cadherin vector (h) were introduced into RGCs by *in utero* electroporation at E12.5. The E13.5 cortices were immunostained for E-cadherin (red). Yellow arrowheads show mClover3^+^ RGCs with apical process extending from the soma to the ventricular wall. Data are shown as mean ± SEM. Unpaired two-tailed Student’s t-test; *p<0.05. Scale bars, 100 µm in (a-c,f); 50 µm in (e,g,h); 20 µm in (d).

RGCs adhere to the apical surface by adherens junctions composed of N-cadherin, E-cadherin, and ZO-1. Although ZO-1 and N-cadherin were observed at the apical surface as they were in the littermate control, the expression of E-cadherin at the apical surface declined in *Mboat7* KO mice (Fig. 3e). Actin filaments at the ventricular wall were unaffected in *Mboat7* KO mice. On the other hand, not only E-cadherin expression but also N-cadherin and ZO-1 expressions at the apical surface decreased, and the disruption of the ventricular zone was observed in the E13.5 cortex of *Mboat7* KO mice (Fig. 3f). These results suggest that the decrease of E-cadherin expression at the apical surface resulted in the detachment of RGCs from the apical surface as a result of *Mboat7* depletion.

### E-cadherin localization at the apical surface is perturbed in LPLAT11-deficient mice

We asked whether the reduced expression of E-cadherin at the apical surface in *Mboat7* KO mice was due to its decreased total expression or its impaired intracellular localization. To this end, mClover3 and E-cadherin expression vectors were introduced into RGCs by *in utero* electroporation at E12.5, and the E13.5 cortex was analyzed. When transfected only with mClover3, the soma of mClover3-positive RGCs exhibited an elongated shape, and some apical processes marked with mClover3 were observed in *Mboat7* heterozygous mice (Fig. 3g). In contrast, the soma showed a rounded shape and the apical process was hardly observed in *Mboat7* KO mice, which supports the idea that the apical processes of RGCs were lost in *Mboat7* KO mice. On the other hand, when RGCs were cotransfected with the E-cadherin expression vector, E-cadherin expression at the apical surface increased and the number of RGC processes at the apical surface also increased in *Mboat7* heterozygous mice (Fig. 3h). However, in *Mboat7* KO mice, E-cadherin was not increased at the apical surface but instead accumulated in the perinuclear region, and the apical processes were not increased, indicating that E-cadherin overexpression *per se* could not rescue the reduced expression of E-cadherin at the apical surface in *Mboat7* KO mice. These results imply that E-cadherin localization at the apical surface is perturbed by LPLAT11 depletion.

### The Golgi apparatus is fragmented in the cortex of LPLAT11-deficient mice

The Golgi apparatus plays a pivotal role in E-cadherin trafficking^37–39^. In RGCs, the Golgi apparatus localizes only to the apical process, and the Golgi apparatus extends as nuclei move upward in cell cycle progression^40^. Immunostaining of E12.5 cortex for GM130, a marker for the Golgi apparatus, revealed that the Golgi apparatus was extended in *Mboat7* heterozygous mice, whereas it was not extended and instead fragmented in *Mboat7* KO mice (Fig. 4a). Measurement of the lengths of the Golgi apparatus at the upper ventricular zone showed that they were shorter in *Mboat7* KO mice (Fig. 4b). However, the Golgi apparatus in E11.5 cortices of *Mboat7* KO mice was not fragmented (Fig. 4c,d). We also examined the morphology of the Golgi apparatus in the E12 cortices, a half-day earlier than E12.5. Sox2^+^; p-H3^+^ RGCs located at the apical surface, revealing that RGCs do not detach from the apical surface in the *Mboat7* KO mice cortices at E12 (Fig. 4e). On the other hand, fragmentation of the Golgi apparatus was observed in the E12 cortices of *Mboat7* KO mice (Fig. 4f,g), suggesting that fragmentation of the Golgi apparatus is the initial phenotype of *Mboat7* KO mice.

**Fig. 4.**
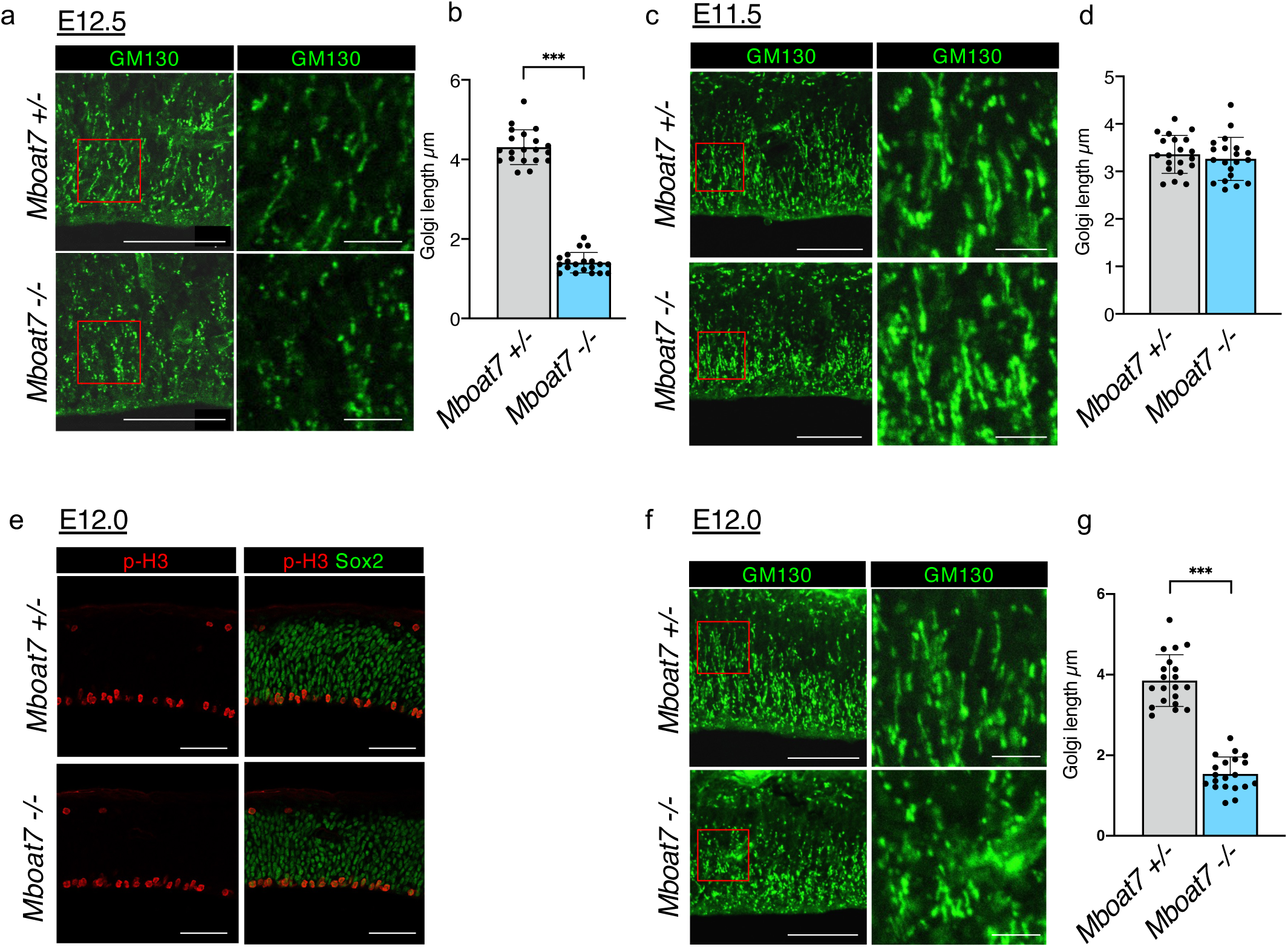
The Golgi apparatus is fragmented from E12 LPLAT11-deficient mice cortex. (a) Immunofluorescence staining for GM130 in the cortices of *Mboat7* heterozygous and KO mice at E12.5. Right panel shows enlarged figure of the red frame. (b) Measurement of the length of the GM130^+^ Golgi apparatus in the ventricular zone (20 cells for each genotype). (c) Immunofluorescence staining for GM130 in the cortices of *Mboat7* heterozygous and KO mice at E11.5. Right panel shows enlarged figure of the red frame. (d) Measurement of the length of the GM130^+^ Golgi apparatus in the ventricular zone (20 cells for each genotype). (e) Double staining for p-H3 and Sox2 in the cortices of *Mboat7* heterozygous and KO mice at E12.0. (f) Immunofluorescence staining for GM130 in the cortices of *Mboat7* heterozygous and KO mice at E12.0. Right panel shows enlarged figure of the red frame. (g) Measurement of the length of the GM130^+^ Golgi apparatus in the ventricular zone (20 cells for each genotype). Data are shown as mean ± SEM. Unpaired two-tailed Student’s t-test; ***p<0.001. Scale bars, 10 µm (enlarged figure in (a,c,f)) and 50 µm (others).

To investigate whether *Mboat7* KO mice showed any abnormality before E11.5, we evaluated the global gene expression patterns in the cortices of *Mboat7* heterozygous mice and KO mice at E11.5 using RNA-sequencing. No significant differences in gene expression were observed (Supplementary Fig. 4), suggesting that *Mboat7* KO mice develop normally until E11.5.

### The total amount of PI(4,5)P_2_ is decreased in LPLAT11-deficient mice cortex

Since fragmentation of the Golgi apparatus is associated with defects in phosphoinositide (PIP) metabolisms^41^, we next measured PI and PIPs in E11.5-E13.5 cortices by using liquid chromatography-mass spectrometry (LC-MS) and supercritical fluid chromatography-mass spectrometry (SFC-MS)^42^, respectively. In *Mboat7* KO mice, PI with arachidonic acid (38:4 PI), which is the major species in *Mboat7* heterozygous mice, significantly decreased, and instead, PI with monounsaturated fatty acid (34:1, 34:2, 36:1, 36:2 PI) and PI with docosapentaenoic acid (DPA) or docosahexaenoic acid (DHA) (38:6, 40:5, 40:6, 40:7 PI) increased as previously reported (Supplementary Fig. 5a-c)^21, 22^. PI4P and PI(4,5)P_2_ also showed similar changes in the molecular species as PI (Supplementary Fig. 5d-i). Interestingly, the total amount of PI(4,5)P_2_ started to decrease at E12.5 in *Mboat7* KO mice, whereas the total amounts of PI and PI4P were unchanged (Fig. 5a-c). This is because the decrease of arachidonic acid-containing PI(4,5)P_2_ species was not fully compensated by the increase of other PI(4,5)P_2_ species (Supplementary Fig. 5h,i). Similar results were obtained by imaging MS analyses in E13.5 cortices (Fig. 5d-f). In *Mboat7* KO mice, arachidonic acid-containing PI and PIP decreased, whereas monounsaturated fatty acid-and DHA-containing PI and PIP increased (Fig. 5d,e). Although arachidonic acid-containing PIP_2_ also decreased, only a subtle increase was observed in monounsaturated fatty acid-and DHA-containing PIP_2_ (Fig. 5f). The composition of molecular species of phosphatidylserine (PS), phosphatidylcholine (PC), and phosphatidylethanolamine (PE) showed little or no change in the cortices of E11.5-E13.5 *Mboat7* KO mice compared with littermate control (Supplementary Fig. 6a-i). Moreover, PI(4,5)P_2_ staining of E13.5 cortices also revealed decreased PI(4,5)P_2_ in the cortex of *Mboat7* KO mice, especially at the apical surface (Fig. 5g).

**Fig. 5.**
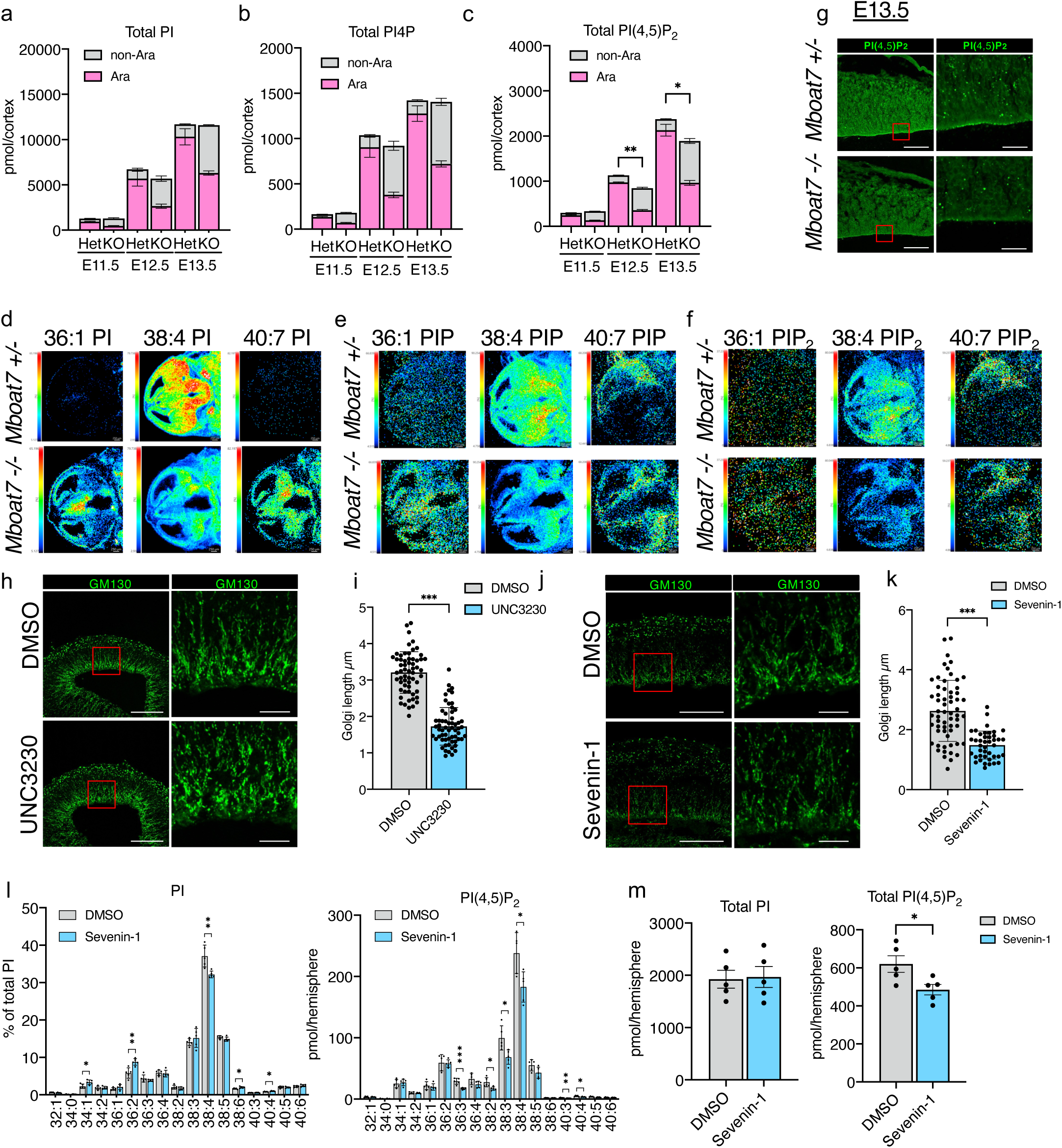
The total amount of PI(4,5)P_2_ decreases in LPLAT11-deficient mice cortex from E12.5. (a) Measurement of total PI in the cortices in *Mboat7* heterozygous and KO mice at E11.5-E13.5 using LC-MS/MS-based method (n=3-5 mice for each genotype). Peak area are normalized by the area of internal standard (25:0 PI). Ara and non-Ara indicate arachidonic acid-containing and non-arachidonic acid-containing species, respectively. (b,c) Measurement of total PI4P (b) and PI(4,5)P_2_ (c) in the cortices in *Mboat7* heterozygous and KO mice at E11.5-E13.5 using SFC-MS/MS-based method (n=3-7 mice for each genotype). Peak area are normalized by the area of internal standard (37:4 PI4P or 37:4 PI(4,5)P_2_). (d-f) Imaging MS analysis of PI (d), PIP (e) and PIP_2_ (f) at E13.5 cortices of *Mboat7* heterozygous and KO mice. (g) Immunofluorescence staining for PI(4,5)P_2_ in the cortices of *Mboat7* heterozygous and KO mice at E13.5. Right panel shows enlarged figure of the red frame. (h) Immunofluorescence staining for GM130 in a cultured E12.5 cortical hemisphere. The Golgi apparatus is fragmented in the hemispheres treated with PIP5K1γ inhibitor, UNC3230. (i) Measurement of the length of the GM130^+^ Golgi apparatus in the ventricular zone (60 cells for each group) in (h). (j) Immunofluorescence staining for GM130 in a cultured E12.5 cortical hemisphere. The Golgi apparatus is fragmented in the hemispheres treated with LPLAT11 inhibitor, Sevenin-1. (k) Measurement of the length of the GM130^+^ Golgi apparatus in the ventricular zone (60 cells for each group) in (j). (l) PI and PI(4,5)P_2_ molecular species in cultured E12.5 hemispheres treated with Sevenin-1 (n=5). (m) Total PI and PI(4,5)P_2_ in cultured E12.5 hemispheres treated with Sevenin-1 (n=5). Data are shown as mean ± SEM. Unpaired two-tailed Student’s t-test; *p<0.05, **p<0.01 and ***p<0.001. Scale bars, 20 µm (enlarged figure in (g,h,j)) and 100 µm (others).

To further examine whether PI(4,5)P_2_ is needed for the proper morphology of the Golgi apparatus in RGCs, we treated E12.5 hemispheres with UNC3230, a PIPKIγ inhibitor in the hemisphere culture. UNC3230 treatment caused fragmentation of the Golgi apparatus, as observed in *Mboat7* KO mice (Fig. 5h,i). We also treated E12.5 *Mboat7* heterozygous hemispheres with Sevenin-1, an LPLAT11 inhibitor^43^. Sevenin-1 treatment caused fragmentation of the Golgi apparatus (Fig. 5j,k), suggesting the importance of LPLAT11 activity. SFC-MS analysis confirmed that Sevenin-1 decreased the amount of PI with arachidonic acid (Fig. 5l) and the total amount of PI(4,5)P_2_ (Fig. 5m).

Together, these results indicate that impaired remodeling to arachidonic acid-containing PI by LPLAT11 depletion results in the decrease of PI(4,5)P_2_ and causes fragmentation of the Golgi apparatus.

### LPLAT11 expression in the cortex determines the integrity of RGCs

To confirm that the phenotype of *Mboat7* KO cortices results from LPLAT11 deficiency in the cerebral cortex, we generated telencephalon-specific *Mboat7* KO mice (hereafter, *Mboat7*-Emx1-KO) by crossing *Mboat7^fl/fl^* mice with Emx1-Cre transgenic mice, in which Cre expression starts at E10.5 in the dorsal telencephalon^44^. The number of cleaved caspase-3-positive cells and c-Jun phosphorylation increased in E13.5 and E14.5 cortices of *Mboat7*-Emx1-KO mice but not as much as in *Mboat7* KO mice (Fig. 6a,b). Moreover, whereas p-H3-positive RGCs were located at the ventricular wall at E12.5, some p-H3-positive RGCs were located apart from the ventricular zone from E13.5 cortices of *Mboat7*-Emx1-KO mice (Fig. 6c). Taken together, these results indicate that telencephalon-specific LPLAT11 deficiency caused the apical detachment, an increase in c-Jun phosphorylation, and apoptosis of RGCs.

**Fig. 6.**
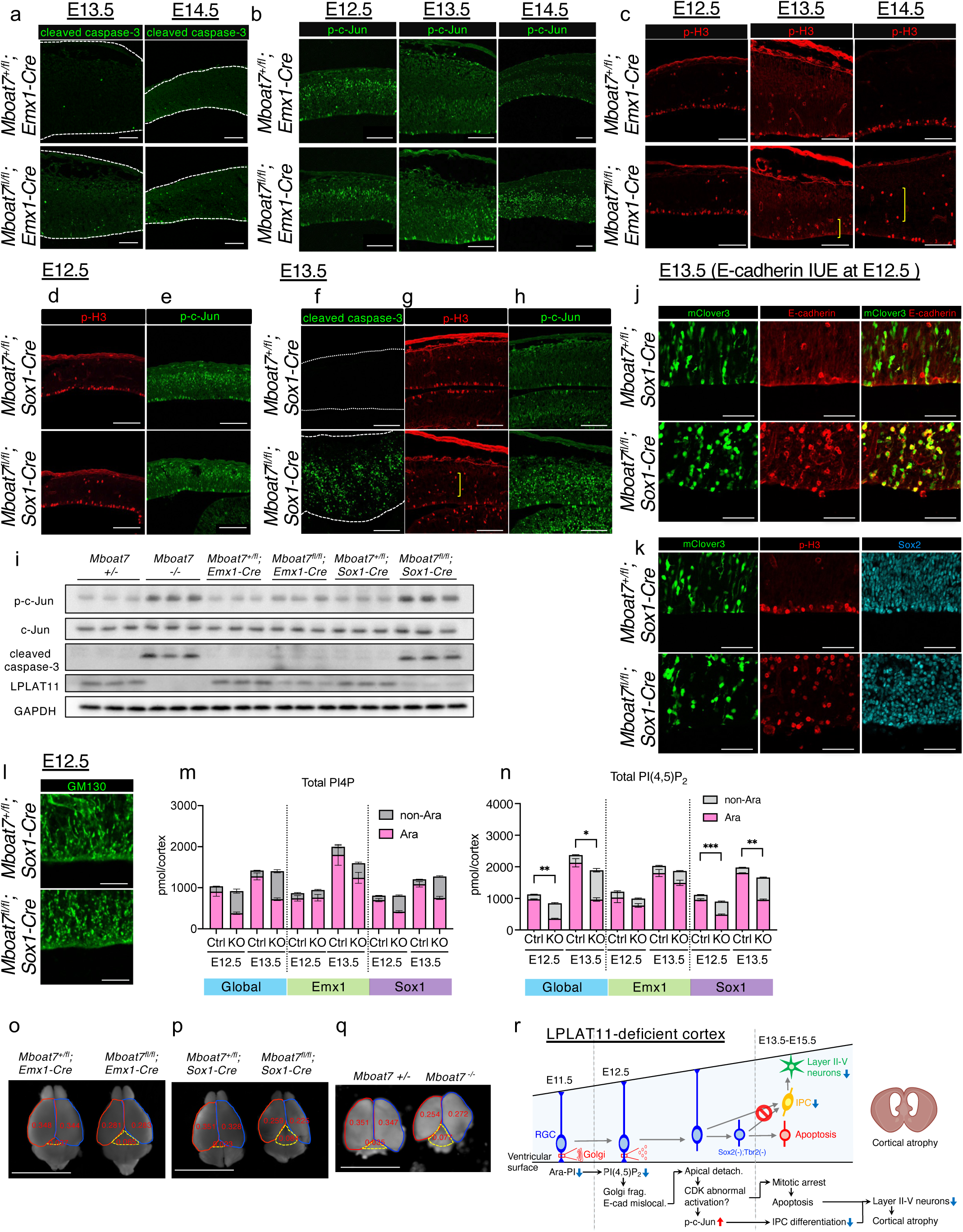
Neural-specific LPLAT11-deficient mice mimic the phenotype of LPLAT11-deficient mice. (a-c) Immunostaining for cleaved caspase-3 (a) in E13.5 and E14.5 cortices, p-c-Jun (b) and p-H3 (c) in E12.5-E14.5 cortices of telencephalon-specific (Emx1-Cre) *Mboat7* heterozygous (*Mboat7*+/fl; Emx1-Cre) and KO mice (*Mboat7*fl/fl; Emx1-Cre). Yellow brackets show RGCs detached from ventricular wall. Dotted lines show the apical and the basal surfaces of the cortex. (d,e) Immunostaining for p-H3 (d) and p-c-Jun (e) in E12.5 cortices of neural-specific (Sox1-Cre) *Mboat7* heterozygous (*Mboat7*+/fl; Sox1-Cre) and KO mice (*Mboat7*fl/fl; Sox1-Cre). (f-h) Immunostaining for cleaved caspase-3 (f), p-H3 (g) and p-c-Jun (h) in E13.5 cortices of neural-specific *Mboat7* heterozygous and KO mice. Dotted lines show the apical and the basal surfaces of the cortex. Yellow bracket shows M phase cells apart from the ventricular wall. (i) Western blots of p-c-Jun, c-Jun, cleaved caspase-3, LPLAT11 in the cortices of global *Mboat7* heterozygous and KO, telencephalon-specific *Mboat7* heterozygous and KO, and neural-specific *Mboat7* heterozygous and KO mice at E13.5 (n=3 mice for each genotype). GAPDH is used as a loading control. (j,k) pCAGGS-mClover3 and pCAGGS-E-cadherin vector were introduced into RGCs of neural-specific *Mboat7* heterozygous and KO mice by *in utero* electroporation at E12.5. The E13.5 cortices were immunostained for E-cadherin (j, red), p-H3 (k, red) and Sox2 (k, cyan). (l) Immunostaining for GM130 in E12.5 cortices of neural-specific *Mboat7* heterozygous and KO mice. (m,n) Quantification of total amounts of PI4P (m) and PI(4,5)P_2_ (n) in E11.5-E13.5 cortices of *Mboat7* heterozygous and KO, telencephalon-specific *Mboat7* heterozygous and KO, and neural-specific *Mboat7* heterozygous and KO mice using SFC-MS/MS based method (n=3-7 mice for each genotype). Ara and non-Ara indicate arachidonic acid-containing and non-arachidonic acid-containing species, respectively. (o-q) Telencephalon-specific *Mboat7* heterozygous and KO mice (o), neural-specific *Mboat7* heterozygous and KO mice (p) or global *Mboat7* heterozygous and KO mice (q) were sacrificed at 3-week-old. Red and blue lines show the outline of left and right cortices. Dotted lines show the outline of midbrain. Areas of cortices and midbrain (cm^2^) are shown in red figures. (r) A schematic diagram of the mechanism for cortical atrophy in *Mboat7* KO mice. Data are shown as mean ± SEM. Unpaired two-tailed Student’s t-test; *p<0.05, **p<0.01, and ***p<0.001. Scale bars, 100 µm in (a-h); 50 µm in (j,k); 20 µm in (l); 1 cm in (o-q).

In *Mboat7*-Emx1-KO mice, the change in the fatty acyl composition of PI and PIPs was delayed compared to *Mboat7* KO mice (Supplementary Fig. 7a,c,e,g) probably because Emx1-Cre drives the expression of Cre recombinase only from E10.5^44^. Therefore, we generated neural-specific *Mboat7* KO mice (hereafter, *Mboat7*-Sox1-KO) by crossing *Mboat7^fl/fl^* mice with Sox1-Cre transgenic mice, which drives the expression of Cre recombinase before E8.5 in the neuroepithelium^45,46^. Detachment of RGCs from the ventricular wall and increased c-Jun phosphorylation were not observed in the E12.5 cortices of *Mboat7*-Sox1-KO (Fig. 6d,e). On the other hand, E13.5 cortices of *Mboat7*-Sox1-KO showed detachment from the ventricular wall, increased c-Jun phosphorylation, and apoptosis to the same extent as those of *Mboat7* KO mice (Fig. 6f-i). Also, *Mboat7*-Sox1-KO showed fragmentation of the Golgi apparatus at E12.5 when the detachment of RGCs from the ventricular wall was not observed (Fig. 6l,d), which is consistent with the phenotype observed in E12 cortices of *Mboat7* KO mice (Fig. 4e). Moreover, E-cadherin overexpression failed to rescue the detachment of RGCs from the apical ventricular wall (Fig. 6j,k). In the *Mboat7*-Sox1-KO cortices, a significant change in the fatty acyl composition of PI and PIPs was observed (Supplementary Fig. 7b,d,f,h). The total amount of PI(4,5)P_2_ started to decrease at E12.5 in the *Mboat7*-Sox1-KO cortices as well as in the *Mboat7* KO cortices (Fig. 6m,n). We also examined whether cortical atrophy was observed in *Mboat7*-Emx1-KO and *Mboat7*-Sox1-KO mice. Cortical atrophy was significant in *Mboat7*-Sox1-KO mice as well as *Mboat7* KO mice, but less in *Mboat7*-Emx1-KO mice (Fig.6o-q). Taken together, these results indicate that *Mboat7*-Sox1-KO mice mimicked almost all the phenotypes observed in the brain of *Mboat7* KO mice.

Notably, *Mboat7*-Sox1-KO mice died within a month after birth (Table 1), indicating that LPLAT11 deficiency in the neuroepithelium is responsible for the postnatal lethality of *Mboat7* KO mice^21^.

**Table 1.** Neural-specific *Mboat7* KO mice die within a month after birth.

| Time | Genotype | *Mboat7*<sup>+/fl</sup> | *Mboat7*<sup>fl/fl</sup> | *Mboat7*<sup>+/fl</sup>; *Sox1*-Cre | *Mboat7*<sup>fl/fl</sup>; *Sox1*-Cre |
| --- | --- | --- | --- | --- | --- |
| 1 week | Number of animals<br>% | 5<br>25.0 | 5<br>25.0 | 7<br>35.0 | 3<br>15.0 |
| 2 week | Number of animals<br>% | 8<br>42.1 | 4<br>21.1 | 6<br>31.6 | 1<br>5.2 |
| 3 week | Number of animals<br>% | 24<br>38.1 | 19<br>30.2 | 20<br>31.7 | 0<br>0.0* |
| Time | Genotype | *Mboat7*<sup>+/fl</sup> | *Mboat7*<sup>fl/fl</sup> | *Mboat7*<sup>+/fl</sup>; *Emx1*-Cre | *Mboat7*<sup>fl/fl</sup>; *Emx1*-Cre |
| 3 week | Number of animals<br>% | 24<br>21.8 | 32<br>29.1 | 34<br>30.9 | 20<br>18.2 |

Genotypes of litters from intercrosses of *Mboat7*<sup>+/fl</sup>; *Sox1*-Cre or *Mboat7*<sup>+/fl</sup>; *Emx1*-Cre with *Mboat7*<sup>fl/fl</sup> mice. DNA was extracted from the ear of each mouse and subjected to PCR analysis to determine the genotype.
The *p* values were calculated using the chi-square test. \**p* = 0.000069

## Discussion

Polyunsaturated fatty acids, such as arachidonic acid and DHA, play a crucial role in embryonic brain development. Although most PUFAs are present in the form bound to phospholipids, little is known about how PUFA-containing phospholipids affect brain development. In this study, we revealed that LPLAT11, the main enzyme that incorporates arachidonic acid into PI, is essential for maintaining the integrity of RGCs in the embryonic mouse brain (Fig. 6r). This is in sharp contrast to the fact that mice deficient in LPLAT12/LPCAT3 (encoded by *Mboat5*), which is a major enzyme responsible for introducing arachidonic acid into PC, PE, and PS, show no significant brain phenotype^47,48^. Thus, the present study suggests arachidonic acid plays a role in brain development by acting as a fatty acyl chain of PI.

RGCs are highly polarized along their apico-basal axis. The architecture and integrity of RGCs at the apical end-feet are maintained by adherens junctions composed of N-cadherin and E-cadherin. We demonstrated that the expressions of E-cadherin and N-cadherin at the ventricular wall in LPLAT11-deficient mice started to decrease at E12.5 and E13.5, respectively. Although adherens junctions disappear as RGCs differentiate into neurons or IPCs, untimely loss of adherens junctions leads to apoptosis. For example, N-cadherin-deficient mice show aberrant accumulation of p-H3-positive cells and increased apoptosis in the cerebral cortex^49,50^. Consistent with this, LPLAT11-deficient mice showed massive apoptosis in the E13.5-E15.5 cortex. Neurons generated during E13.5-E15.5 form layers II-V of the cerebral cortex, in which atrophy was observed in LPLAT11-deficient mice. During that period, there was no decrease in the proliferation of RGCs or failure of these cells to differentiate into neurons. These results suggest that apoptosis leads to cortical atrophy in LPLAT11-deficient mice.

We found that the apoptotic cells expressed neither Sox2, Tbr2, nor Ctip2, marker proteins for RGCs, IPCs, and neurons, respectively. Cells undergoing apoptosis can become negative for any marker proteins^51^. However, some p-H3-positive cells that did not undergo apoptosis were also negative for Sox2 and Tbr2 in the LPLAT11-deficient cortex, suggesting that the loss of marker protein expressions occurred prior to apoptosis. In general, as RGCs differentiate into immature IPCs, the expression of RGC markers decreases, while the expression of IPC markers increases^52^. Phosphorylation of c-Jun, which inhibits Tbr2 expression in immune cells^31^, was upregulated, and differentiation of RGCs into IPCs was impaired in LPLAT11-deficient mice. These results suggest that aberrant c-Jun activation suppresses the expression of Tbr2 and differentiation into IPCs, resulting in Sox2/Tbr2-negative cells that eventually undergo apoptosis in LPLAT11-deficient mice.

Another characteristic of the apoptotic cells is that they were not labeled with BrdU, which indicates that cells whose entry into the S-phase is impaired undergo apoptosis. Moreover, p-H3-positive RGCs accumulated in the E13.5 cortex of LPLAT11-deficient mice. These results suggest that LPLAT11-deficient RGCs have undergone mitotic arrest. Consistently, aberrant mitotic processes, including dysregulation of CDKs, lead to a mitotic catastrophe that ultimately induces apoptosis^32^. In fact, we demonstrated that a CDK inhibitor reduced the number of p-H3-positive RGCs and suppressed c-Jun phosphorylation and apoptosis in the cortex of LPLAT1-deficient mice. It is noteworthy that loss of adherens junction in RGCs by N-cadherin KO or expression of dominant negative N-cadherin leads to mitotic catastrophe, as evidenced by the increased number of p-H3-positive cells, cells negative for Sox2 and Tbr2, and massive apoptotic cells^49,50^. Thus, these findings suggest that decreased expression of E-cadherin and N-cadherin in RGCs causes mitotic catastrophe due to aberrant CDK activation, resulting in c-Jun phosphorylation, suppression of Tbr2 expression, and induction of apoptosis in the cortex of LPLAT11-deficient mice.

*In utero* electroporation of the E-cadherin expression vector failed to restore the expression of E-cadherin at the apical surface in the LPLAT11-deficient cortex. Instead, transfected E-cadherin localized at the perinuclear region, suggesting that E-cadherin transport to the apical membrane is disturbed in LPLAT11-deficient mice. The Golgi apparatus of RGCs has a unique organization: Golgi apparatus stacks reside only in the apical process and are oriented with their *cis*-to-*trans* axis perpendicular to the apical-basal axis of the cell, which contributes to the transport of apical membrane proteins. We found that the Golgi apparatus of RGCs was fragmented in the E12 cortex of LPLAT11-deficient mice, where no other cellular abnormalities were noted. This indicates that fragmentation of the Golgi apparatus is the initial cellular defect of LPLAT11 deficiency. Moreover, a decreased level of arachidonic acid-containing PI by LPLAT11 deficiency led to a reduction of the PI(4,5)P_2_ level in the E12.5 cortex. PI(4,5)P_2_ is critical for the trafficking of apical membrane proteins^53,54^. It has also been shown that PI(4,5)P_2_ is localized not only at the plasma membrane but at the Golgi apparatus, where PI(4,5)P_2_ participates in the structural organization of the Golgi apparatus^41^. Thus, the PI(4,5)P_2_ reduction in the RGCs may impair the transport of E-cadherin by affecting the trafficking machinery at the plasma membrane and/or the organization of the Golgi apparatus.

Arachidonic acid-containing PI and PIPs were decreased while PI and PIPs with fatty acids other than arachidonic acid were increased in E11.5 cortices of LPLAT11-deficient mice. However, the decrease in the amount of PI(4,5)P_2_ in the cortex of LPLAT11-deficient mice was not observed until E12.5, suggesting that PI(4,5)P_2_ synthesis is perturbed starting at E12.5. PI(4,5)P_2_ is generated from PI4P by PI4P 5-kinases (PIPKIs). Among the three PIPKI isoforms (α, β, γ), PIPKIγ is the dominant isoform in the brain since PIPKIγ KO mice show a decreased amount of PI(4,5)P_2_ in the brain while PIPKIα KO, PIPKIβ KO, and PIPKIα/β DKO mice do not^55^. Expression of PIPKIγ in the brain is detected in the E12 cortex and increases with embryonic development^55^. PIPKIγ has been reported to prefer arachidonic acid-containing PI4P as a substrate over saturated fatty acid-containing PI4P^56^. Therefore, PIPKIγ, whose expression in the brain starts at E12.5 brain, may fail to produce a sufficient amount of PI(4,5)P_2_ in LPLAT11-deficient cortex, where the level of arachidonic acid in PI4P is reduced. Alternatively, a reduced level of PI with arachidonic acid may affect the function of phosphatidylinositol transfer proteins in PIPs metabolism at the Golgi apparatus since mice deficient in phosphatidylinositol transfer proteins (*Pitpna/Pitpnb* DKO) show a phenotype similar to LPLAT11-deficient mice^57^.

Patients harboring null mutations in *MBOAT7* showed polymicrogyria, which is thought to be caused by wrong aberrant neuronal migration. We previously demonstrated that neuronal migration was delayed, and neurons were scattered in the cortex of LPLAT11-deficient mice^21^. Since radial fibers serve as scaffolds for radial neuronal migration, the disarrangement of radial fibers is likely the main cause of inappropriate neuronal migration in LPLAT11-deficient mice. Thus, the disorganized arrangement of radial fiber may also occur in the cortex of the patients. The layer II-V neurons, which were decreased in LPLAT11-deficient mice, are callosal projection neurons connecting the two hemispheres in the cerebral cortex. Since defects in these neurons are associated with intellectual disability, loss of these neurons may underlie intellectual disability in patients harboring null mutations in *MBOAT7*.

In conclusion, the present study demonstrated the role of LPLAT11-mediated PI remodeling in the integrity of RGCs, which provides a mechanistic insight into the importance of arachidonic acid in embryonic brain development.

## Methods

### Mice

LPLAT11-deficient mice were generated previously^17^. Emx1-Cre mice (No.005628) were purchased from the Jackson Laboratory. Sox1-Cre mice were provided by RIKEN (No. CDB0525K, https://large.riken.jp/distribution/mutant-list.html). These transgenic mice were crossed with *Mboat7^flox/flox^* mice. All mice were housed in climate-controlled (23 °C) pathogen-free facilities with a 12 h light-dark cycle, with free access to standard chow (CE2; CLEA Japan) and water. All animal experiments were performed in accordance with protocols approved by the Animal Committees of the University of Tokyo in accordance with the Standards Relating to the Care and Management of Experimental Animals in Japan. For timed pregnancy, the morning of the detection of the vaginal plug was designated as embryonic day E0.5.

### Antibodies

Antibodies to cleaved caspase-3 (#9661), Sox2 (#3728S), phospho-Histone H3 (#9706S), and phospho-c-Jun (#9261) were purchased from Cell Signaling Technology, Inc. Antibodies to Sox2 (sc-365964) and c-Jun (sc-1694) were purchased from Santa Cruz Biotechnology. Antibodies to Tbr1 (Ab31940), Ctip2 (Ab18465), Tbr2 (Ab23345), and BrdU (Ab6326) were purchased from Abcam. Antibody to Cux1 (NBP2-13883) was purchased from Novus Biologicals. Antibody to PI(4,5)P_2_ (Z-P045) was purchased from Echelon Biosciences. Antibody to ZO-1 (Z-R1) was purchased from Zymed Laboratories. Antibodies to E-cadherin (610181), N-cadherin (610920), and Nestin (611658) were purchased from BD Transduction Laboratories. Antibodies to GAPDH (6C5), Pax6 (AB2237), and Tbr2 (AB15894) were purchased from Merck Millipore. Antibody to LPLAT11 was generated in our laboratory^21^.

### BrdU Labeling

For BrdU labeling, E12.5 or E13.5 pregnant mice were given intraperitoneal injection with 50 mg/kg BrdU. To investigate differentiation from RGCs into neurons or IPCs, BrdU injections were given 24 h before sacrifice. To label cells in the S/G2 phase, early G1 phase, and late G1 phase, mice were injected with BrdU 2 h, 6 h, and 14 h before harvesting.

### Immunohistochemistry

For immunofluorescence staining, brains were postfixed with ice-cold 4% (wt/vol) paraformaldehyde (PFA) at 4 °C for 2-3 h, equilibrated with 15% (w/v) sucrose in PBS at 4 °C overnight, 30% (w/v) sucrose in PBS at 4 °C overnight, and frozen in OCT (Tissue TEK). Coronal sections (14 µm thickness) were exposed to TBST buffer (25 mM Tris-HCl, pH7.5, 140 mM NaCl, 0.1% Triton X-100) for 30 min (permeabilization) and 3% BSA-TBST (blocking) for 1 h at room temperature and incubated overnight at 4 °C with primary antibodies in blocking buffer. For staining with the antibodies to Cux1, Ctip2, Tbr1, Tbr2, and BrdU, we performed antigen retrieval by autoclave treatment of sections with 10 mM citrate buffer (pH6.0) for 10 min at 105 °C. For staining with the antibody to PI(4,5)P_2_, TBS containing 0.01% saponin was used for permeabilization, and 3% BSA-TBS containing 0.01% saponin was used for blocking. Sections were incubated with Alexa-Fluor-labeled secondary antibodies and DAPI in a blocking buffer for 2 h at room temperature.

Fluorescence images were obtained with a laser confocal microscope (Leica TCS-SP5 and SP8). Alexa-Fluor-labeled secondary antibodies and DAPI were obtained from Merck Millipore.

### *Ex vivo* culture of cortical hemisphere

Embryos were dissected at E12.5, and two cerebral cortical hemispheres were separated as described previously^35^ with some modification. Briefly, the brains of embryos were dissected in individual 35 mm dishes containing culture medium (Opti-MEM I with 20 mM glucose, 55 µM 2-mercaptoethanol, and 1% penicillin/streptomycin). Separate hemispheres were transferred to a new 35m dish containing culture medium using a P1000 Pipetman with a cut pipette tip and kept on ice until every dissection was completed. Hemispheres were transferred to an individual well of a 12-well plate containing 2 ml of culture medium. Plates were then placed on a nutator inside a 5% CO_2_ incubator for 24 h.

For CDK inhibitor treatment, Indirubin-3’-monoxime (I3M) was obtained from Cayman Chemical Company and was added at 1 µM final concentration in the culture medium. For LPLAT11 inhibitor treatment, N-(2-(Bicyclo[2.2.1]heptanyl)ethyl)-4-(2-(N-methylsulfamoyl)phenyl)piperazine-1-carboxamide (Sevenin-1)^43^ was obtained from Enamine Company and was dissolved in DMSO. The inhibitor was added at 10 µM final concentration in the culture medium. For PIPKIγ inhibitor treatment, UNC3230 was obtained from TargetMol Chemicals Inc. and was dissolved in DMSO. The inhibitor was added at 0.25 µM final concentration in the culture medium.

### CDK inhibitor injection into the lateral ventricle

I3M was diluted to 10 µM in PBS, mixed with the same amount of CFSE containing fastgreen, and injected into the lateral ventricle of each littermate at E11.5. PBS containing 0.1% DMSO was used as a control. The uterine horn was placed back into the abdominal cavity to allow embryos to continue development. The next day, the embryos were collected for immunohistochemical analysis of the brain.

### *In utero* electroporation

Introduction of plasmid DNA into the neuroepithelial cells of mouse embryo *in utero* was performed as described^58^. Plasmid DNA for pCAGGS-mclover3 (1 µg/µl) + pCAGGS-empty vector (2 µg/µl) or pCAGGS-mclover3 (1 µg/µl) + pCAGGS-mouse E-cadherin (2 µg/µl) were injected into the lateral ventricle of each littermate at E12.5. Electrodes were placed flanking the equivalent ventricular region of each embryo, covered with a drop of PBS, and pulsed 8 times at 40 V for 50 ms, separated by intervals of 950 ms with an electroporator (CUY21E; NEPA Gene). The uterine horn was placed back into the abdominal cavity to allow the embryo to continue development. The next day, the embryos were collected for immunohistochemical analysis of the brain.

### Western blotting

Cortices from E12.5 to E18.5 were isolated in ice-cold PBS including 8 mM NaF, 12 mM beta-glycerophosphate, 1 mM Na_3_VO_4_, 1.2 mM Na_2_MoO_4_, 5 µM Cantharidin, and 2 mM Imidazole and collected in lysis buffer (62.5 mM Tris-HCl pH6.8, 10% Glycerol, 1% SDS) and sonicated. The lysate was incubated at room temperature for 30 min and then centrifuged at 15,000 g for 30 min at room temperature, and the supernatants were collected. The protein concentration was determined by the BCA assay (Pierce). Proteins were separated by SDS-PAGE and transferred to PVDF membranes (Millipore). The membranes were blocked with 5 % skimmed milk in TTBS buffer (10 mM Tris-HCl, pH 7.4, 150 mM NaCl, 0.05% Tween 20) for 1 h at room temperature and then incubated with primary antibodies overnight at 4°C. On the next day, the membranes were incubated with horseradish peroxidase-conjugated anti-mouse, anti-rabbit, or anti-rat antibody (GE Healthcare). Proteins were detected by enhanced chemiluminescence (ECL western blotting detection system, GE Healthcare) using the ImageQuant LAS 4000 system (GE Healthcare).

### Lipid extraction

Lipid extractions were conducted by the method of Bligh and Dyer^59^. The extracted solutions were dried up with a centrifugal evaporator, dissolved in chloroform/methanol (2/1, v/v), and stored at −20 °C.

For preparation for SFC-MS samples, cortices from E11.5 to E13.5 were isolated in ice-cold DMEM and immediately collected into a safe-lock poly-propylene tube (2ml) with 750 µl of ice-cold quench mix and sonicated. 170 µl of H_2_O was mixed and stored at −80 °C before lipid extraction. Lipid extraction was performed based on procedures by J Clark *et al.*^60^. The single-phase sample (a mixture of 170 µl of an aqueous sample, 10 ng of internal standards [17:0/20:4 PI, 17:0/20:4 PI4P, and 17:0/20:4 PI(4,5)P_2_], and 750 µl of quench mix) were mixed with 725 µl of CHCl_3_ and 170 µl of 2 M HCl, followed by vortex-mixing and centrifugation (1,500 g, 5 min at room temperature). The lower organic phase was collected into a new 2 ml safe-lock poly-propylene tube and mixed with 708 µl of pre-derivatization wash solution, followed by vortex-mixing and centrifugation (1,500 g, 3 min at room temperature). The lower phase was collected into another fresh tube and subjected to derivatization, as described below.

### Derivatization of extracted lipids

Derivatization of lipids was performed in a fume hood with adequate personal safety equipment as follows, based on procedures described by J Clark *et al.*^60^. 50 µl trimethylsilyl diazomethane in hexane (2 M solution; Sigma-Aldrich) was added to the lipid extracts (∼1 ml), and after leaving to stand 10 min at room temperature, the reaction was quenched with 6 µl of acetic acid. Next, 700 µl post-derivatization wash solution was added to the mixture, followed by centrifugation (1,500 g, 3 min). The resultant lower phase was collected and rewashed with a 700 µl post-derivatization wash solution. Samples were temporarily stored at −80 °C. Ninety µl of MeOH and 10 µl of H_2_O were added to the final collected lower phase. The samples were dried up with a centrifugal evaporator. Samples were dissolved in 80 µl MeOH, sonicated for 1 min, and 20 µl H_2_O was added. After centrifugation at 15,000 g for 30 min at 4 °C, the supernatants were collected. To avoid degradation, the samples were stored at −80 °C until use.

### MS analysis of phospholipids

LC/ESI-MS–based lipidomics analyses were performed on a Shimadzu Nexera UPLC system (Shimadzu) coupled with a QTRAP 4500 hybrid triple quadrupole linear ion trap mass spectrometer (AB SCIEX) as previously described^44^. Chromatographic separation was performed on a SeQuant ZIC-HILIC PEEK coated column (250 mm × 2.1 mm, 1.8 µm; Millipore) maintained at 50 °C using mobile phase A (water/acetonitrile (95/5, v/v) containing 10 mM ammonium acetate) and mobile phase B (water/acetonitrile (50/50, v/v) containing 20 mM ammonium acetate) in a gradient program (0–22 min: 0% B→40% B; 22–25 min: 40% B→40% B; 25–30 min: 0% B) with a flow rate of 0.3 ml/min.

### SFC-MS analysis of phosphoinositides

For the detection of phospholipids, SFC-MS/MS-based lipidomics analyses were performed on a Shimadzu Nexera UC/s system (Shimadzu) coupled with a QTRAP 4500 hybrid triple quadrupole linear ion trap mass spectrometer (AB SCIEX). Lipids extracted were injected by an autosampler; typically, 10 µl of the sample was applied. Chromatographic separation was performed on an ULTRON AF-HILIC-CD (250 mm x 2.1 mm, 5.0 µm; Shinwa Chemical Industries) maintained at 10 °C using immersion cooler Neo cool drip BE201F (Yamato Scientific) with mobile phase A [supercritical carbon dioxide (SCCO_2_)] and mobile phase B [water/methanol (2.5/97.5, v/v) containing 0.1% (v/v) formic acid] in a gradient program (0-16 min: 5% B→20% B; 16.01-18 min: 40% B; 18.01-22 min: 5% B) with a flow rate of 1.5 ml/min. The instrument parameters of QTRAP4500 for positive ion mode were as follows: curtain gas, 20 psi; ionspray voltage, 4500 V; temperature, 300 °C; ion source gas 1, 18 psi; ion source gas 2, 20 psi.

### MS imaging

E13.5 brains were embedded in 2% carboxymethylcellulose and frozen in liquid nitrogen. The brain sections (10 µm) were washed by submerging in 50 mM of ammonium formate for 5 s. MALDI matrix *p*-NA was applied to sections as previously described^61^. Briefly, *p*-NA was dissolved in 80% EtOH containing 150 mM ammonium formate. The section was sprayed with a SunCollect automated sprayer at an air pressure of 0.3 MPa and a 15 μL/min flow rate. First, the brain sections were sprayed with 10 layers of 1 mg/mL *p*-NA, and then 25 layers of 10 mg/ mL *p*-NA. MALDI-MSI analyses were performed using Fourier transform orbital trapping MS (QExactive, Thermo Fisher Scientific, San Jose, CA) connected to a MALDI laser system (AP-SMALDI5, TransMIT, Giessen, Germany) in full-scan mode at a mass resolution of 140,000. The laser was manually focused. The laser power was set at 27° attenuator setting to yield optimal results. Ion images were reconstructed using IMAGEREVEAL^TM^ MS (Shimadzu, Kyoto, Japan).

### RNA-seq

E11.5 cortex was dissected in ice-cold DMEM, and total RNA was isolated with RNeasy Mini Kit (Qiagen) and RNase-Free DNase Set (79254, Qiagen) following the manufacturer’s instructions. Sequencing was performed at Rhelixa, Inc., Japan, using the Illumina NovaSeq 6000 platform.

### Statistical analysis

Data were presented as means ± s.e.m. values, and the unpaired Student’s *t-test* was applied to determine significant differences between the two samples.

## Data availability

The datasets generated and/or analyzed during the current study are available in the manuscript or the Supplementary Information and are available from the corresponding author upon request.

## Acknowledgments

We thank S. Nishikawa (RIKEN) for providing *Sox1-Cre* mice. This work was supported by grants from the Japan Society for the Promotion of Science KAKENHI, grant numbers 23K14343 (to Y. I.), 17H06164 and 17H06418 (to H. A.); the Japan Agency for Medical Research and Development, grant numbers JP21gm1210013 (to N. K.), and JP21gm0010004h9905 and JP22ck0106533h0003 (to J. A.); Japan Science and Technology Agency Moonshot R&D Program, grant numbers JPMJMS2023-11 (to J. A.) and JPMJMS2023-15 (to J. A.).

## Contributions

Y.I. and N.Kono designed and conceptualized the study. Y.I. Y.K. T.I., and N.Kuwayama were responsible for experiments and data generation. H.A. Y.G., and J.A. supervised the research. Y.I. and N.Kono wrote the original and revised drafts of the paper. All authors read and approved the paper.

## Ethics declarations

Competing interests

The authors declare no competing interests.

---

**Supplementary Fig. 1.**
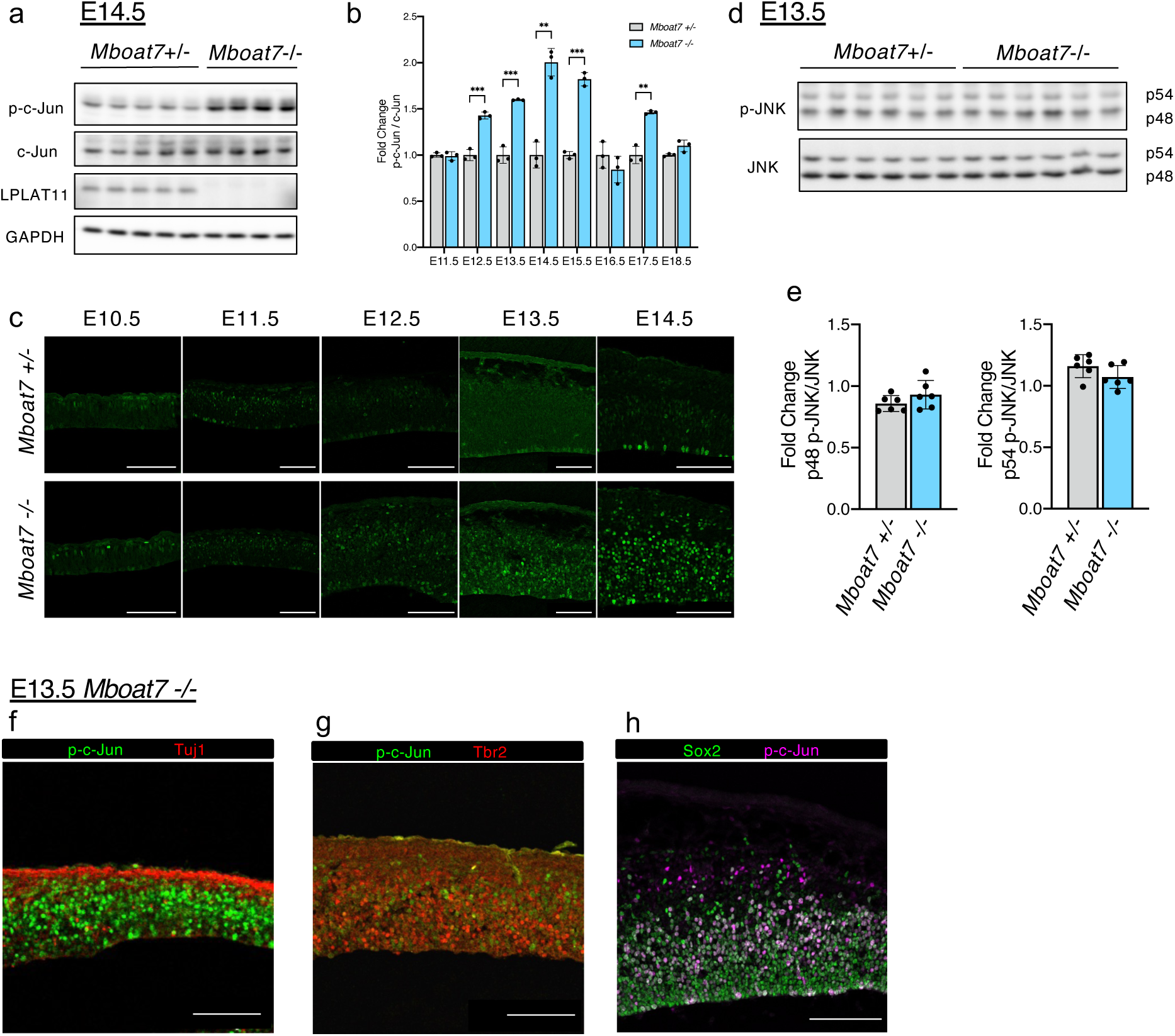
Phosphorylation of c-Jun is upregulated in LPLAT11-deficient RGCs. (a) Western blots of p-c-Jun, c-Jun and LPLAT11 in E14.5 cortices of *Mboat7* heterozygous and KO mice. GAPDH is used as a loading control. (b) Evaluation of c-Jun phosphorylation in the cortices of *Mboat7* heterozygous and KO mice from E11.5-E18.5 (n=3 mice for each genotype). Fold change against *Mboat7* heterozygous mice is shown. (c) Immunofluorescence staining for p-c-Jun in the cortices of E10.5-E14.5 *Mboat7* heterozygous and KO mice. (d) Western blots of p-JNK and JNK in E13.5 cortices of *Mboat7* heterozygous and KO mice. (e) Evaluation of p48 and p54 JNK phosphorylation in E13.5 cortices of *Mboat7* heterozygous and KO mice (n=6 mice for each genotype). (f-h) Double staining for p-c-Jun and Tuj1 (f), Tbr2 (g) and Sox2 (h) in the cortices of *Mboat7* KO mice at E13.5. Scale bars, 100 µm. Data are shown as mean ± SEM. Unpaired two-tailed Student’s t-test; **p<0.01 and ***p<0.001.

**Supplementary Fig. 2.**
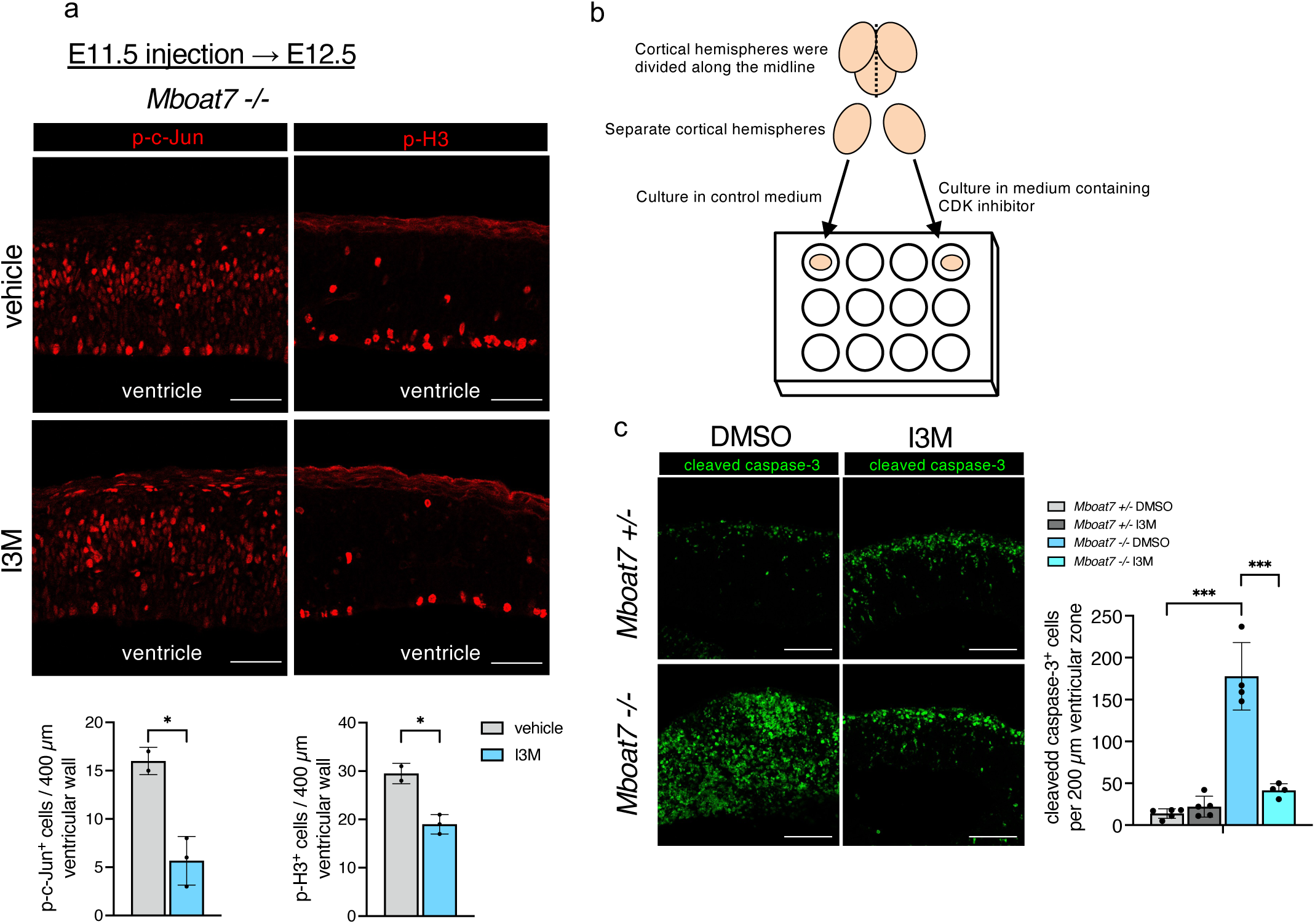
LPLAT11 deficiency shows increased apoptosis by aberrant activation of CDK. (a) CDK inhibitor (Indirubin 3’-monoxime, I3M) was administered into the ventricle of *Mboat7* KO mice at E11.5. The E12.5 cortices were immunostained for p-H3. Graphs show quantitative analysis of cells highly positive for p-c-Jun (left) and positive for p-H3 per 400 µm ventricular wall (n=2,3 mice for each group). Data are shown as mean ± SEM. Unpaired two-tailed Student’s t-test; *p<0.05. (b) Illustration of a cortical hemisphere culture. Cortical hemispheres were separated along the midline and were cultured in medium containing DMSO or CDK inhibitor (I3M), respectively. (c) Hemispheres cultured in medium with DMSO or 1µM CDK inhibitor (I3M) were immunostained for cleaved caspase-3. Cleaved caspase-3^+^ cells per 200 µm ventricular zone were shown (n=4,5 mice for each group). Data are shown as mean ± SEM. One-way ANOVA with Tukey’s post hoc test; ***p<0.001. Scale bars, 50 µm in (a); 100 µm in (c).

**Supplementary Fig. 3.**
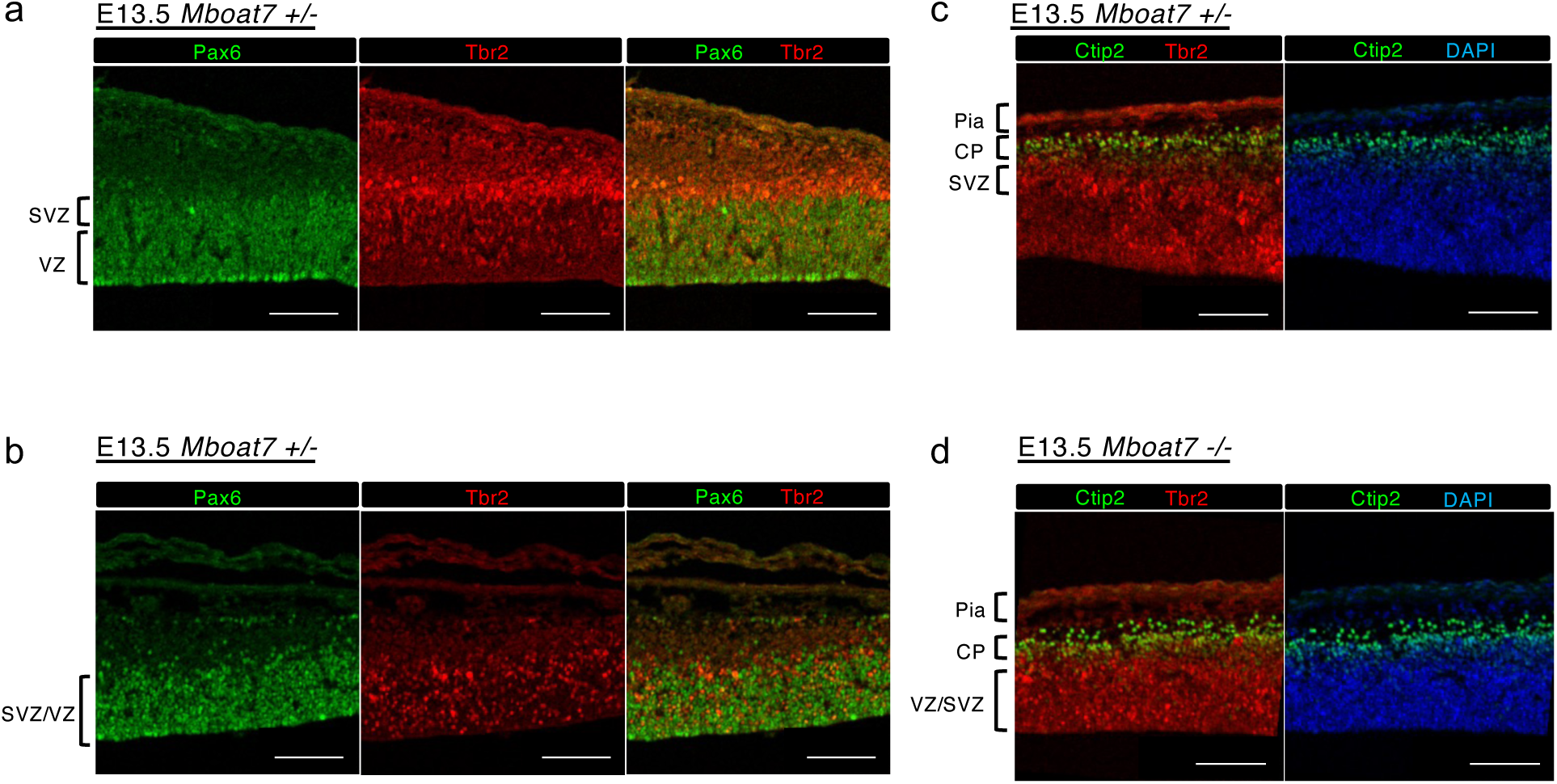
RGCs and IPCs are scattered in LPLAT11-deficient mice. (a,b) Expression of Pax6 (green) and Tbr2 (red) in E13.5 cortices of *Mboat7* heterozygous and KO mice. (c,d) Expression of Ctip2 (green) and Tbr2 (red) in E13.5 cortices of *Mboat7* heterozygous and KO mice. Nuclei were stained with DAPI. Pia, pial surface; CP, cortical plate; VZ, ventricular zone; SVZ, subventricular zone. Scale bars, 100 µm

**Supplementary Fig. 4.**
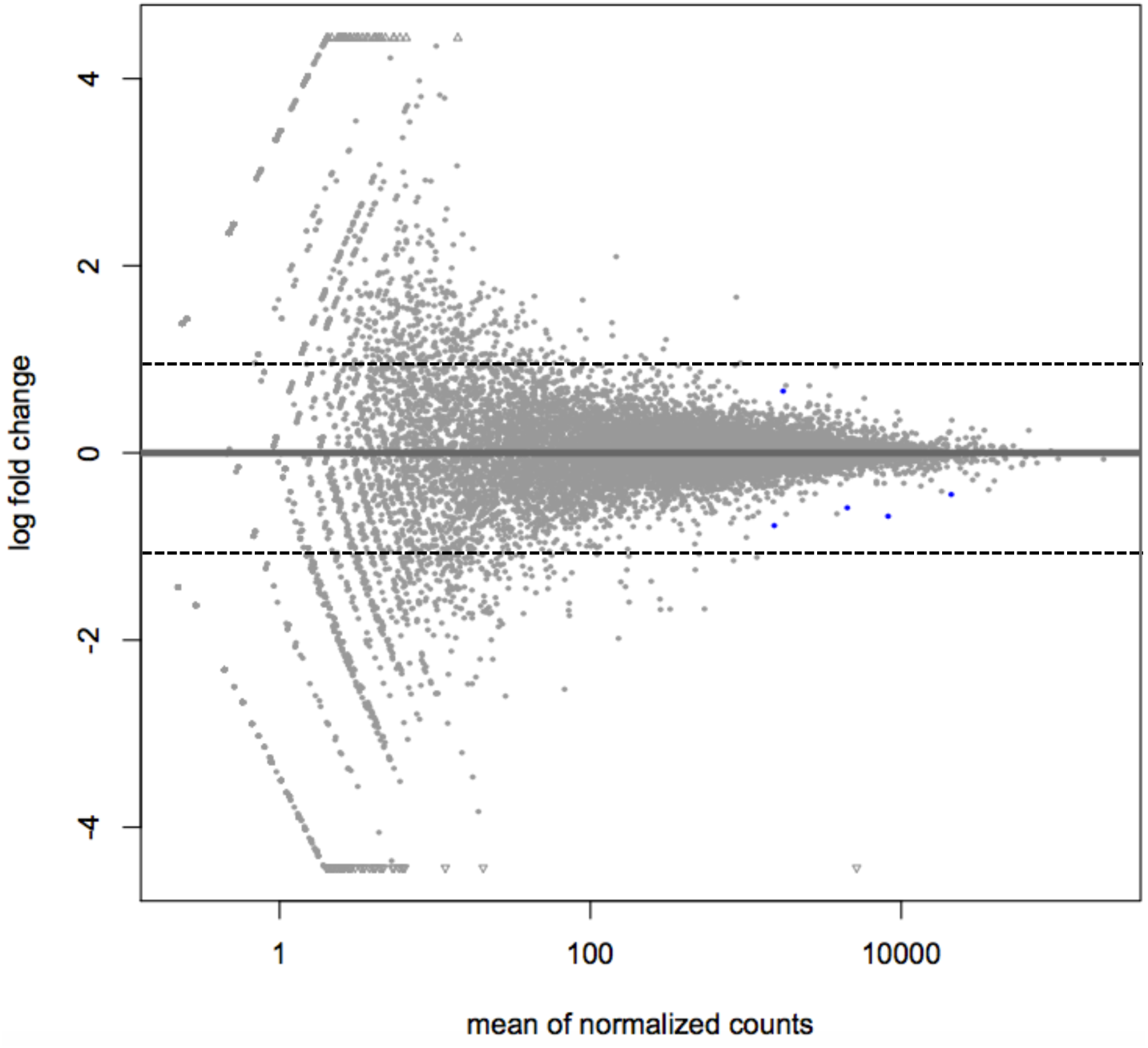
RNA-seq analysis of E11.5 cortices. Plot of normalized mean versus log2 fold change for the contrast *Mboat7* heterozygous mice versus *Mboat7* KO mice. Dotted lines show log2 fold change −1, 1, respectively. Blue dots show statistically significantly changed genes in *Mboat7* KO mice.

**Supplementary Fig. 5.**
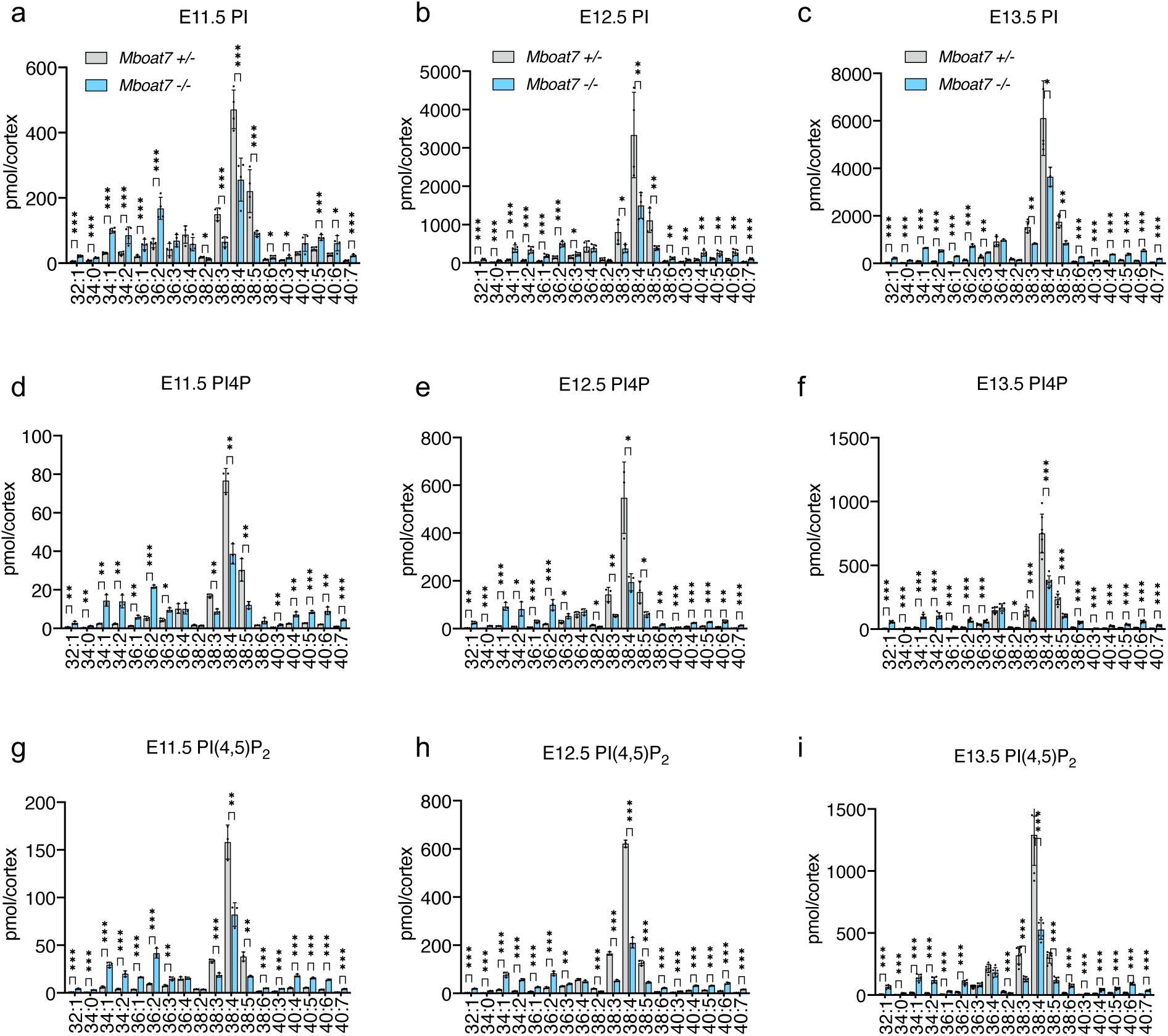
Fatty acid composition of PI, PI4P and PI(4,5)P_2_ in the cortices of E11.5-E13.5 of LPLAT11-deficient mice. (a-c) LC-MS/MS analysis of PI at E11.5 (a), E12.5 (b) and E13.5 (c) cortices of *Mboat7* heterozygous and KO mice (n=3-5 mice for each genotype). Peak area are normalized by the area of internal standard (25:0 PI). (d-f) SFC-MS/MS analysis of PI4P molecular species at E11.5 (d), E12.5 (e) and E13.5 (f) cortices of *Mboat7* heterozygous and KO mice (n=3-5 mice for each genotype). Peak area are normalized by the area of internal standard (37:4 PI4P). (g-i) SFC-MS/MS analysis of PI(4,5)P_2_ molecular species at E11.5 (g), E12.5 (h), and E13.5 (i) cortices of *Mboat7* heterozygous and KO mice (n=3-5 mice for each genotype). Peak area are normalized by the area of internal standard (37:4 PI(4,5)P_2_). Data are shown as mean ± SEM. Unpaired two-tailed Student’s t-test; *p<0.05, **p<0.01, and ***p<0.001.

**Supplementary Fig. 6.**
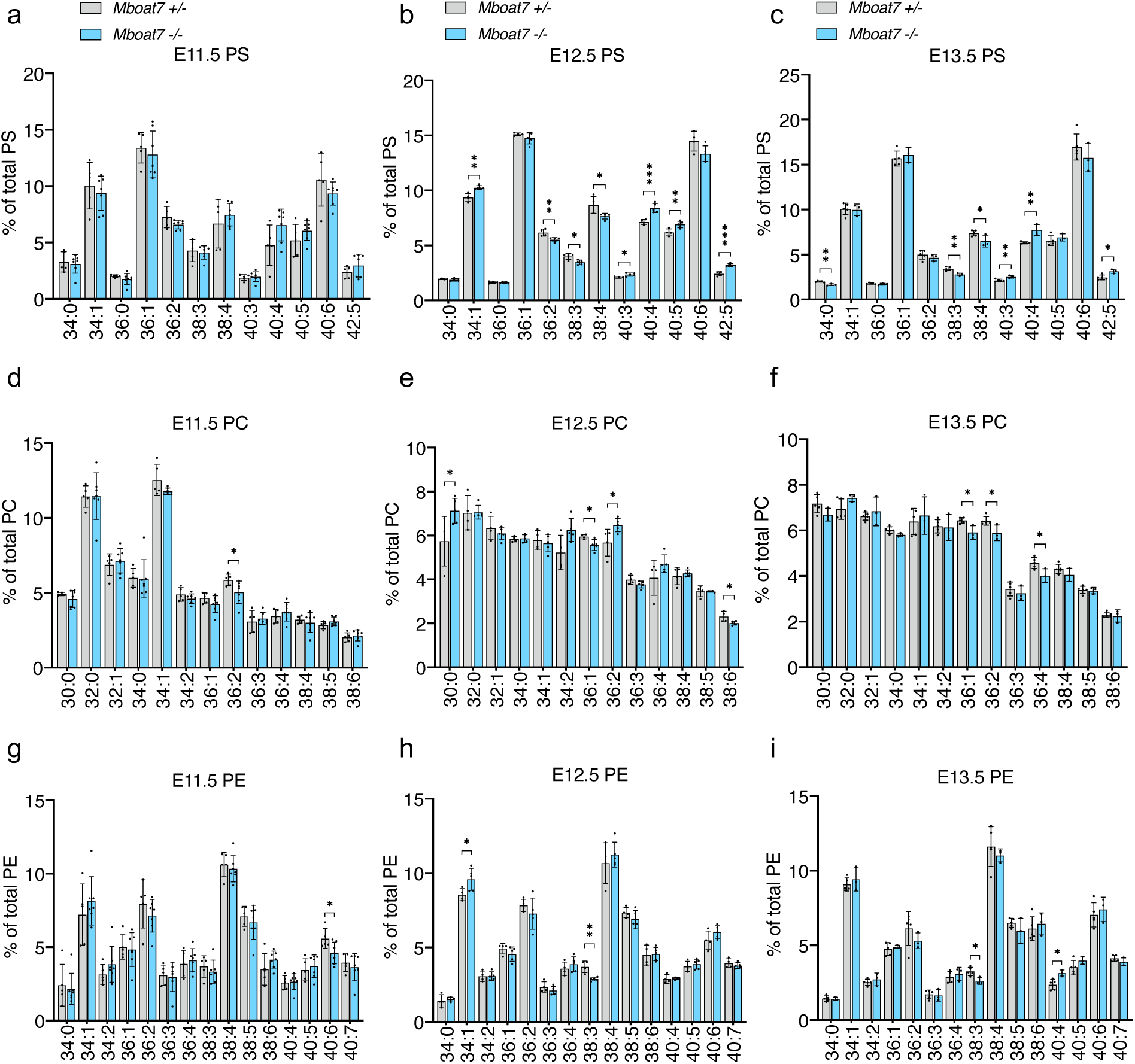
LC-MS/MS analysis of phospholipid species in the cortices of E11.5-E13.5 mice. (a-c) LC-MS/MS analysis of PS at E11.5 (a), E12.5 (b) and E13.5 (c) cortices of *Mboat7* heterozygous and KO mice (n=3-5 mice for each genotype). Peak area are normalized by the area of internal standard (25:0 PS). (d-f) LC-MS/MS analysis of PC at E11.5 (d), E12.5 (e) and E13.5 (f) cortices of *Mboat7* heterozygous and KO mice (n=3-5 mice for each genotype). Peak area are normalized by the area of internal standard (25:0 PC). (g-i) LC-MS/MS analysis of PE at E11.5 (g), E12.5 (h) and E13.5 (i) cortices of *Mboat7* heterozygous and KO mice (n=3-5 mice for each genotype). Peak area are normalized by the area of internal standard (25:0 PE). Data are shown as mean ± SEM. Unpaired two-tailed Student’s t-test; *p<0.05, **p<0.01, and ***p<0.005.

**Supplementary Fig. 7.**
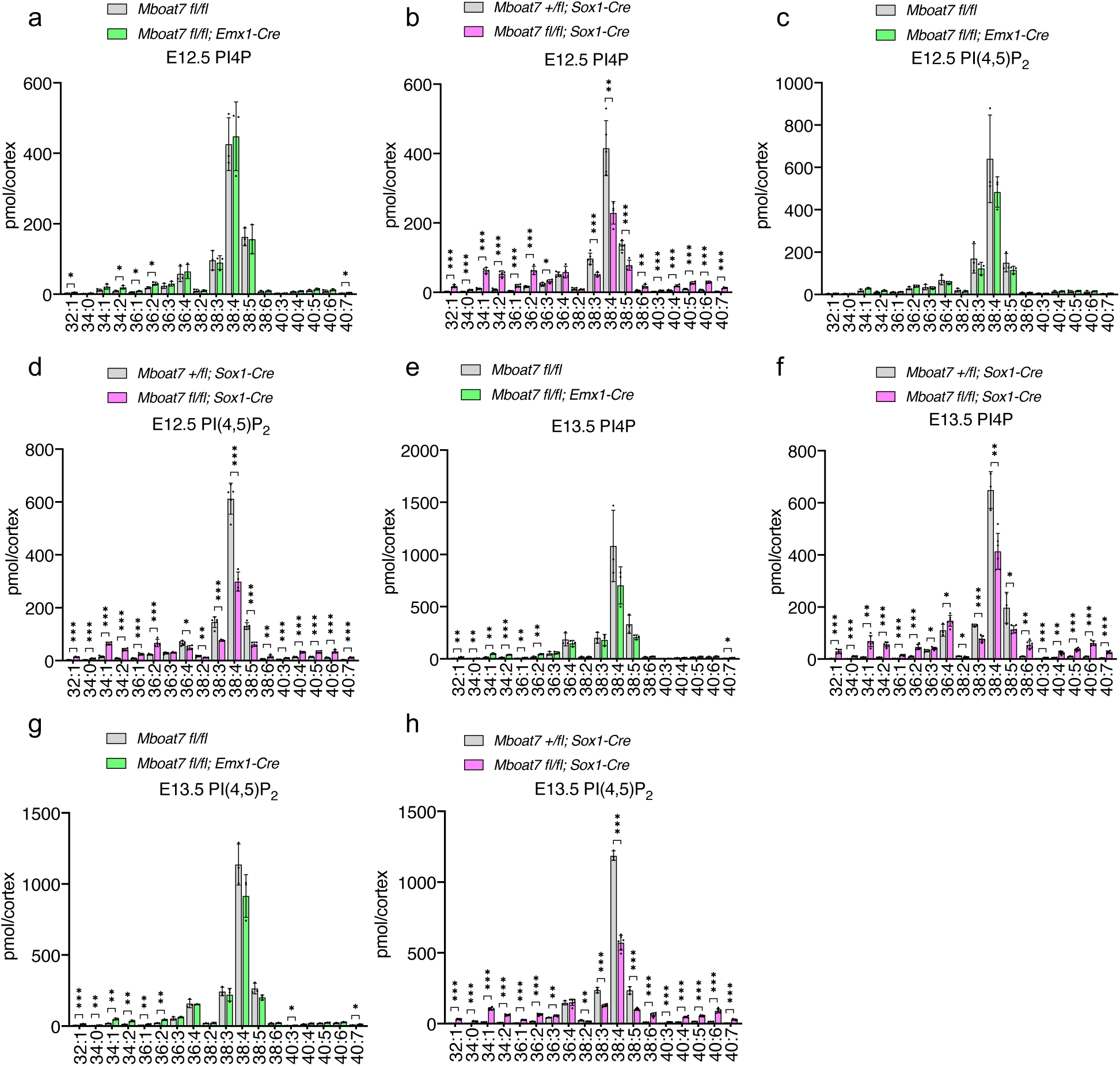
SFC-MS/MS analysis in the cortices of conditional LPLAT11-deficient mice at E12.5 and E13.5. (a,b) SFC-MS/MS analysis of PI4P molecular species in E12.5 cortices of telencephalon-specific (Emx1-Cre) *Mboat7* heterozygous and KO mice (a) or neural-specific (Sox1-Cre) *Mboat7* heterozygous and KO mice (b). (c,d) SFC-MS/MS analysis of PI(4,5)P_2_ molecular species in E12.5 cortices of telencephalon-specific (Emx1-Cre) *Mboat7* heterozygous and KO mice (c) or neural-specific (Sox1-Cre) *Mboat7* heterozygous and KO mice (d). (e,f) SFC-MS/MS analysis of PI4P molecular species in E13.5 cortices of telencephalon-specific (Emx1-Cre) *Mboat7* heterozygous and KO mice (e) or neural-specific (Sox1-Cre) *Mboat7* heterozygous and KO mice (f). (g,h) SFC-MS/MS analysis of PI(4,5)P_2_ molecular species in E13.5 cortices of telencephalon-specific (Emx1-Cre) *Mboat7* heterozygous and KO mice (g) or neural-specific (Sox1-Cre) *Mboat7* heterozygous and KO mice (h). Peak area were normalized by the area of internal standard (37:4 PI4P or 37:4 PI(4,5)P_2_) (n=3-6 mice for each genotype). Data are shown as mean ± SEM. Unpaired two-tailed Student’s t-test; *p<0.05, **p<0.01, and ***p<0.001.

